# Sequential adaptor function of Treacle and MDC1 couples nucleolar reorganization to RNF8-dependent recruitment of HDR factors

**DOI:** 10.1101/2025.04.21.649841

**Authors:** Andrea Haenel, Johannes Leyrer, Polyxenia Stutz, Manuel Stucki

## Abstract

The repair of DNA double-strand breaks in repetitive sequences is challenging because the abundance of potential templates for homology-directed repair (HDR) increases the risk of ectopic recombination and chromosome rearrangements. Relocalization of repair sites in repetitive sequences to a ‘safe location’ prior to RAD51 loading has been observed in various organisms and is thought to limit aberrant recombination events.

Here, we characterize this process in ribosomal DNA (rDNA) repeats, which reside within the nucleoli, specialized nuclear compartments dedicated to ribosome biogenesis. Induction of DSBs in rDNA triggers large-scale mobilization of damaged repeats from the nucleolar interior to the nucleolar periphery, where repair occurs. We show that this process is coordinated by the adaptor proteins Treacle and MDC1, which act sequentially to control nucleolar reorganization and repair factor recruitment.

Treacle promotes rDNA mobilization through its role in nucleolar DNA damage signalling, whereas MDC1 acts downstream of nucleolar segregation to facilitate the recruitment of HDR factors at nucleolar caps. Following establishment of a γH2AX chromatin domain, MDC1-dependent RNF8-mediated chromatin ubiquitylation promotes BRCA1 recruitment via both the RNF168-dependent pathway and the RAP80–ABRAXAS complex. This in turn enables the sequential accumulation of PALB2 and RAD51 at nucleolar caps.

Together, our findings demonstrate that RAD51 loading at rDNA breaks is tightly coupled to nucleolar reorganization and is mediated by dual RNF8-dependent signalling pathways, thereby ensuring that HDR is spatially restricted to the nucleolar periphery and reducing the risk of ectopic recombination between repetitive sequences.

## Introduction

The human genome is constantly threatened by extrinsic and intrinsic DNA damaging agents, generating a myriad of DNA lesions every day that need to be repaired to prevent accumulation of mutations and chromosomal instability. Amongst various DNA lesions, DNA double-strand breaks (DSBs) are particularly toxic because, if not repaired accurately, DSBs can result in loss of genetic information and/or chromosomal rearrangements. To avoid these lethal events, cells have evolved highly sophisticated mechanisms, collectively termed as DNA damage response (DDR), to detect DNA lesions, signal their presence and promote their repair (Jackson and Bartek 2009). The two main pathways families that facilitate the repair of DSBs are the non-homologous end joining (NHEJ) pathways and homology-directed repair (HDR) pathways. While NHEJ represent the predominant mechanism of DSBs repair, which operates throughout the cell cycle, HDR is mainly active in the late S/G2 phase, when identical sister chromatids are present and used as repair templates (Scully et al. 2019; Chapman et al. 2012).

An early cellular response to DSBs is phosphorylation of the histone variant H2AX at its C-terminus by the ATM kinase. The phosphorylated form of H2AX (termed γH2AX) is recognized by the adaptor protein MDC1, which itself mediates the recruitment of several DDR factors into γH2AX-decorated chromatin regions (Jungmichel and Stucki 2010). Among the proteins recruited by MDC1 is the E3 ubiquitin ligase RNF8 that poly-ubiquitylates the linker histone H1, thus triggering the recruitment of several ubiquitin binding factors including RNF168, another E3 ubiquitin ligase that mono-ubiquitylates Lys 13 and 15 of H2A (Stewart et al. 2009; Kolas et al. 2007; Huen et al. 2007; Mailand et al. 2007; Doil et al. 2009; Thorslund et al. 2015). These modifications flag chromatin for subsequent recruitment of the 53BP1/ Shieldin-CST and BRCA1/BARD1 DDR complexes, respectively, which regulate DSB repair pathway choice during the cell cycle (reviewed in (Hustedt and Durocher 2016).

HDR in repetitive DNA sequences is challenging because repeated DNA sequences could potentially be used as homology template, giving rise to non-homologous (ectopic) recombination. The problem is likely worse when the repeats are located on different chromosomes because in that case, ectopic recombination could occur between different chromosomes, which would lead to translocations and chromosome fusions. For the repair of DSB in pericentric heterochromatin, where an abundance of repeated DNA sequences within the heterochromatin domains is likely to promote ectopic recombination, cells have evolved an elegant solution. Studies performed in Drosophila and mouse cells have shown that early repair steps occur inside the heterochromatin domains, but later steps only occur after a striking relocalization of repair sites to the nuclear periphery (in the case of Drosophila cells) or the periphery of the heterochromatin domains (termed chromocenters) in mouse cells (reviewed in (Amaral et al. 2017)).

A similar mechanism appears to have evolved for the rDNA repeats that code for the 45S precursor of the ribosomal RNA. In human cells, several hundred rDNA repeats are unevenly distributed on the short arms of the five acrocentric chromosomes, where, along with adjacent sequences, they form the nucleolar organising regions (NORs). The NORs from different acrocentric chromosomes are located in close proximity to each other within the nucleoli, membrane less subnuclear compartment where ribosome biosynthesis takes place (van Sluis and McStay 2017).

Targeted DSB induction in the rDNA repeats by either CRISPR/Cas9 or expression of the homing endonuclease I-Ppo1 leads to rapid transcriptional shutdown of the RNA Polymerase I (Pol I) holoenzyme and large-scale movement of rDNA and associated proteins from inside the nucleoli to the nucleolar periphery (van Sluis and McStay 2015; Harding et al. 2015; Warmerdam et al. 2016). The nucleolar low complexity protein Treacle is a key regulator of this response. Treacle (TCOF1) is the protein mutated in Treacher Collins syndrome, a developmental disorder primarily attributed to impaired ribosome biogenesis and nucleolar dysfunction during neural crest development. Treacle directly interacts with NBS1 and TOPBP1 and upon rDNA break induction, mediates their recruitment in the nucleoli, where they promote and sustain ATM and ATR signalling (Larsen et al. 2014; Mooser et al. 2020; Korsholm et al. 2019).

This Treacle-mediated mobilization of rDNA breaks resembles the movement of breaks in heterochromatin repeats, which are only repaired once the breaks have been moved out of chromocenteres and/or towards the nuclear envelope (Amaral et al. 2017). Yet, how these events are coordinated in space and time in the nucleoli remains unknown. Here we show that the two adaptors Treacle and MDC1 are involved in the spacially and timely coordination of nucleolar segregation and recruitment of the HDR machinery. We demonstrate that recruitment of BRCA1 and loading of RAD51 are dependent on both the Treacle-NBS1-TOPBP1 complex and the γH2AX-MDC1-RNF8-RNF168 cascade. Thereby, Treacle-NBS1-TOPBP1 acts upstream of γH2AX-MDC1-RNF8-RNF168, thus ensuring that transcriptional repression and nucleolar segregation occurs before the HDR machinery is recruited, which may help to physically separate rDNA repeats located on separate chromosomes before they are repaired by HDR.

## Results

### The Treacle-TOPBP1-NBS1 complex controls the recruitment of HDR factors to rDNA DSBs

We previously showed that the nucleolar adaptor protein Treacle is regulating nucleolar segregation through its ability to directly interact with the two DNA damage response adaptor proteins NBS1 and TOPBP1 and thus recruit them in the nucleoli, which ultimately facilitates ATR activation (Larsen et al. 2014; Mooser et al. 2020). We were next interested if Treacle was also required for the recruitment of the HDR machinery, which was previously shown to accumulate in the nucleolar periphery upon rDNA break induction (van Sluis and McStay 2015; Harding et al. 2015). To study specifically the cellular response to breaks induced in the rDNA repeats, we transfected cells with *in vitro* transcribed and poly-adenylated mRNA of the I-Ppo1 homing endonuclease, which recognizes a sequence within the 28S rDNA region as well as a few sites elsewhere in the human genome. As a negative control, we used a catalytically dead mutant of I-Ppo1 (H98A).

Consistent with previously published work (van Sluis and McStay 2015) we observed that I-Ppo1 WT, but not H98A mRNA transfection leads to rapid nucleolar segregation and recruitment of the HDR factors BRCA1 and RAD51 to the nucleolar periphery (Figure 1A). The two DNA end resection factors CtIP and BRCA1 accumulated inside of the nucleoli within 1 hour after I-Ppo1 mRNA transfection, whereas PALB2 and RAD51 foci were observed in the nucleolar periphery at later time points (2h), indicating a sequential recruitment of HDR factors rather than the assembly of a pre-formed complex (Figure 1–figure supplement 1A,B).

**Figure 1.**
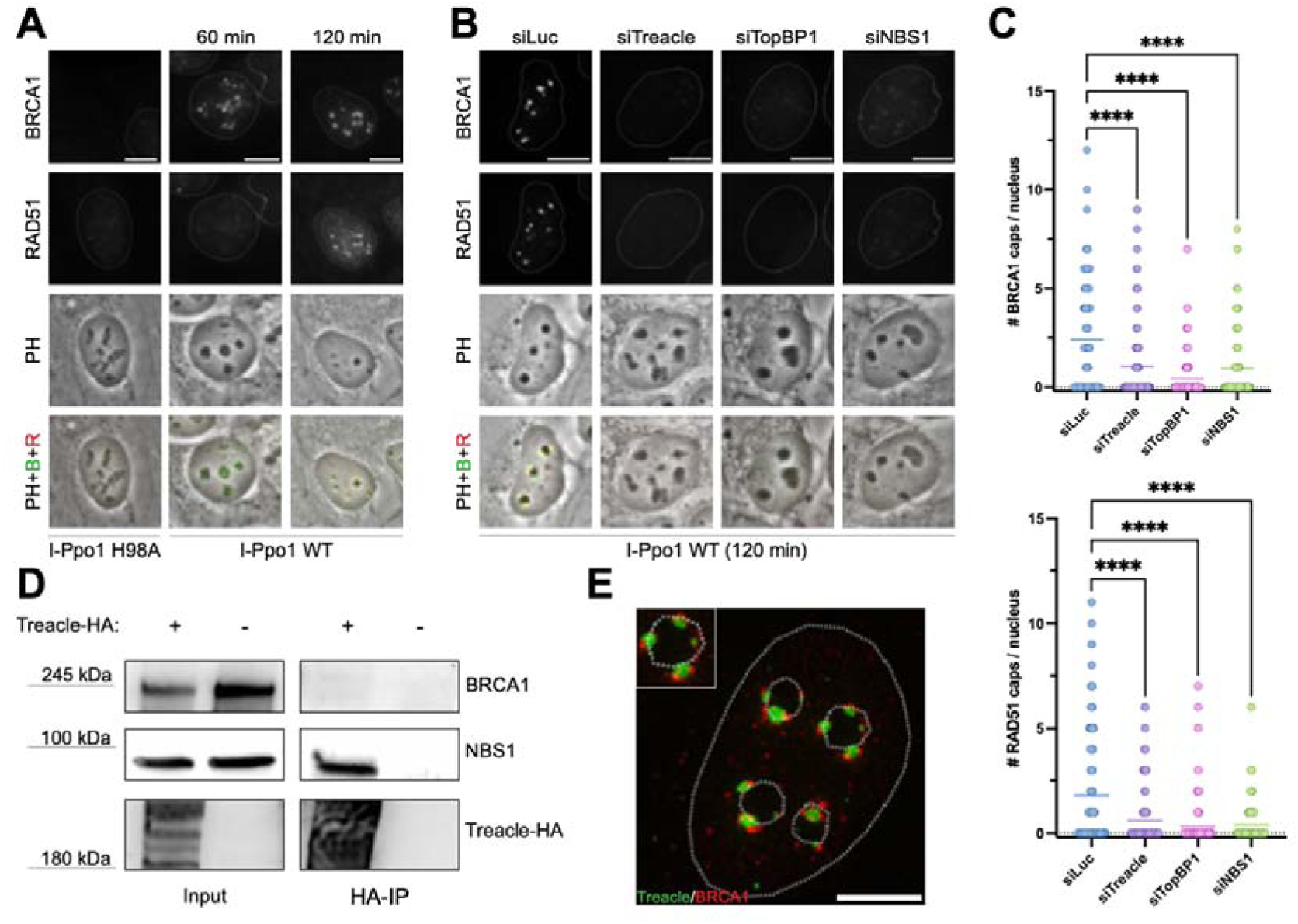
(A) Localisation of BRCA1 and RAD51 after 1h or 2h of I-Ppo1 WT treatment or 2h of I- Ppo1 H98A treatment, respectively, in U2OS cells. (B) Localisation of BRCA1 and RAD51 in siLuc, siTreacle, siTopBP1, and siNBS1 transfected U2OS cells after 2h of I-Ppo1 WT treatment. All scale bars = 10 µm. (C) Quantification of the experiment in (B) showing the number of BRCA1 and RAD51, respectively, nucleolar caps (n= 100 cells). Graphs show a single representative replicate (out of three), and bars represent mean. (D) Co-immunoprecipitation from 293T cells transfected or non-transfected with HA-tagged Treacle and treated with WT I-Ppo1 for 2h. (E) SIM of Treacle and BRCA1 after 2h of I-Ppo1 WT transfection of Treacle-GFP expressing U2OS cells. Scale bar = 5 µm.

Next, we asked if Treacle and its two interaction partners NBS1 and TOPBP1 were required for the recruitment the HDR machinery to broken rDNA repeats. BRCA1 and RAD51 foci within the nucleolar periphery were strongly reduced in Treacle, NBS1 and TOPBP1 depleted cells (Figure 1B and Figure 1–figure supplement 1C).

Given that I-PpoI also induces DNA breaks outside of the rDNA, we next sought to distinguish between nucleolar and non-nucleolar DNA damage responses. To this end, we analyzed the formation of γH2AX-positive nucleolar caps and γH2AX foci over time following I-PpoI expression. While both γH2AX foci and nucleolar caps increased after I-PpoI induction, nucleolar caps appeared rapidly and were prominent at early time points, whereas γH2AX foci accumulated more gradually over time (Figure 1–figure supplement 2A). Importantly, these two classes of structures could be reliably distinguished using a CellProfiler-based image analysis pipeline that segmented a donut-shaped area around the nucleoli (Figure 1–figure supplement 3, see also Methods), which thus allowed us to separate objects in the nucleolar periphery from objects elsewhere in the nucleus (Figure 1–figure supplement 2B). These observations indicate that nucleolar caps represent a spatially and temporally distinct response to rDNA damage and support their use as a specific readout for nucleolar DNA repair events in subsequent experiments.

To quantitatively assess BRCA1 and RAD51 foci specifically around the nucleoli we used our Cell Profiler analysis pipeline’s ability to restrict the identification of focal objects to the nucleolar periphery. This analysis also revealed a significant reduction of BRCA1 and RAD51 foci in the nucleolar periphery area in the absence of Treacle, NBS1 and TOPBP1 (Figure 1C).

Treacle may mediate recruitment of the HDR machinery to the nucleoli directly, e.g. by directly interacting with BRCA1 in a similar manner as it interacts with NBS1 and TOPBP1, respectively. However, we could not detect any interaction between Treacle and BRCA1 by co-immunoprecipitation, whereas interaction with NBS1 was readily detectable (Figure 1D). Moreover, detailed analysis of Treacle and BRCA1 localization in the nucleolar periphery by structured illumination super-resolution microscopy (SIM) revealed largely non-overlapping localization of Treacle and BRCA1 within nucleolar caps, with BRCA1 foci mostly occurring adjacent to Treacle foci (Figure 1E).

Together, these data suggest that Treacle and its two associated DDR factors NBS1 and TOPBP1 may control the accumulation of the HDR factors BRCA1 and RAD51 to sites of rDNA breaks indirectly, most likely via ATR-mediated repression of rRNA transcription and consequential nucleolar segregation.

### Phosphorylation of H2AX and recruitment of MDC1 are essential for the presence of HDR factors at the sites of rDNA DSBs

While SIM microscopy revealed that Treacle and BRCA1 did not co-localized within nucleolar caps, it clearly showed that BRCA1 co-localized with γH2AX and MDC1 signals (Figure 2A,B). We used the 3D rendering feature of the Imaris software packages to reconstruct the obtained SIM images, which provided insight into the detailed structure of nucleolar caps. We observed two separate structures within the nucleolar caps, in which the Treacle signal was limited to the inner sub-structure of nucleolar caps, where it was surrounded by BRCA1 and MDC1 signal that co-localized in the outer shell-like sub-structure of nucleolar caps (Figure 2B).

**Figure 2.**
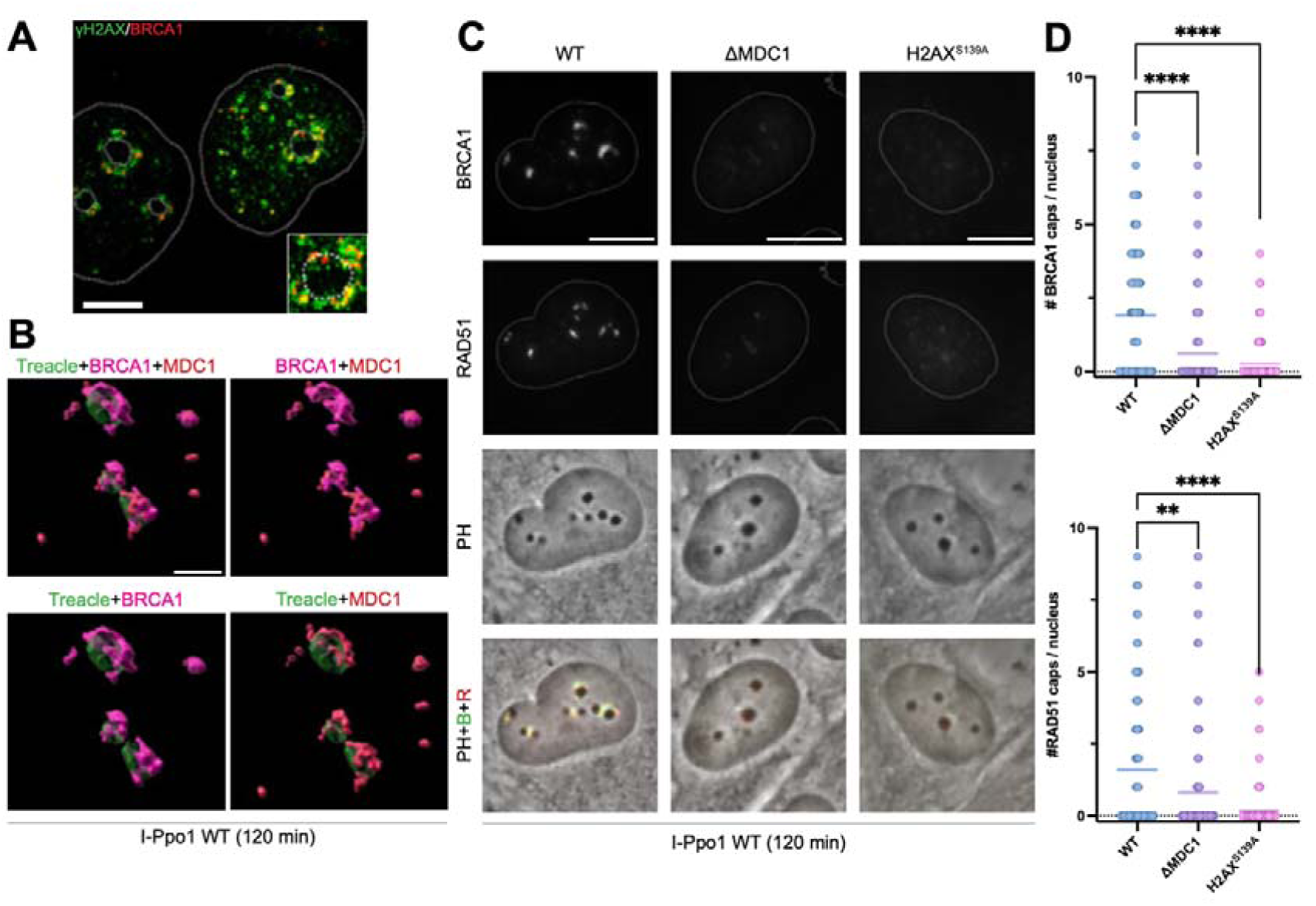
(A) SIM of H2AX and BRCA1 in U2OS cells after 2h of I-Ppo1 WT transfection. Scale bar = 5 μm. (B) SIM of U2OS cells stably expressing Treacle-GFP and treated with I-Ppo1 WT for 2h, stained with BRCA1 and MDC1 antibodies. A reconstruction of nucleolar caps using 3D surface is shown. Scale bar = 1μm. (C) Localisation of BRCA1 and RAD51 in WT, ΔMDC1, and H2AX^S139A^ RPE1 cells 2h after I-Ppo1 WT transfection. All scale bars = 10 µm. (D) Quantification of the experiment in (C) showing the number of BRCA1 (upper panel) and RAD51 (lower panel) caps per cell (n= 90 cells). Graph shows a single replicate (experiment was performed in triplicates) and bars represent mean.

To study the direct involvement of γH2AX and MDC1 in the mobilisation of HR factors to the site of rDNA DSBs, we used RPE1 MDC1 knock-out (ΔMDC1) and bi-allelic RPE1 H2AX S139A knock-in (H2AX^S139A^) cell lines. The accumulation of both BRCA1 and RAD51 in nucleolar caps upon rDNA DSBs was significantly reduced in ΔMDC1 and H2AX^S139A^ cells when compared to the wildtype control cells (Figure 2C-D). To confirm these findings, we used two additional MDC1 knock-out cell lines produced in our laboratory. (Leimbacher et al. 2019). Both U2OS and HeLa ΔMDC1 cells showed significantly reduced recruitment of BRCA1 and RAD51 to the sites of rDNA DSBs (Figure 2–figure supplement 1A.B). This was surprising because previous studies revealed alterations in (but not the abolition of) RAD51 focus formation in H2AX and MDC1 null cells (Scully and Xie 2013; Zhang et al. 2005). We therefore compared BRCA1 and RAD51 DNA damage recruitment after I-Ppo1 transfection to ionizing radiation induced foci (IRIF). Whereas BRCA1 recruitment was defective in U2OS ΔMDC1 cells irrespective of the source of DNA damage, RAD51 recruitment was specifically defective after I-Ppo1, but not after IR-induced break formation (Figure 2–figure supplement 1C).

In summary, these findings suggest that recruitment of the HDR factors to nucleolar caps in response to rDNA breakage is dependent on the γH2AX-MDC1 pathway.

### RNF8-dependent chromatin ubiquitylation pathways mediate BRCA1 and RAD51 recruitment to nucleolar caps

To test which of the regions in MDC1 are required for BRCA1 and RAD51 recruitment to nucleolar caps after rDNA break induction by I-Ppo1, we complemented U2OS ΔMDC1 cells with various MDC1 mutants, including S168A/S196A double mutant previously shown to be defective for TOPBP1 binding (Leimbacher et al. 2019), SDT motif mutant previously shown to be defective for NBS1 binding (Spycher et al. 2008; Melander et al. 2008), TQXF mutant shown to be defective for RNF8 recruitment (Mailand et al. 2007; Kolas et al. 2007; Huen et al. 2007) and PST repeat deletion mutant (DPST) lacking the central repeat region previously shown to be required for γH2AX-independent chromatin interaction (Salguero et al. 2019). All mutants except the RNF8 binding mutant (AQXF) complemented BRCA1 and RAD51 recruitment to nucleolar caps (Figure 3A-C, Figure 3–figure supplement 1), indicating that RNF8 plays a key role in the recruitment of the HDR machinery to the nucleolar periphery. Indeed, RNF8 depletion in U2OS cells stably transfected with an inducible shRNF8 construct (Mailand et al. 2007) resulted in near-complete abrogation of BRCA1 and RAD51 recruitment to nucleolar caps (Figure 4A,B and Figure 4–figure supplement 1A). It was recently shown that BRCA1 is recruited to sites of DSBs by two redundant pathways, both dependent on chromatin ubiquitylation by RNF8 (Sherker et al. 2021). One pathway is also dependent on RNF168, which generates the H2AK13/15ub mark that BARD1 is recognising via its ubiquitin interaction motif (Becker et al. 2021). The second pathway is independent of RNF168 and instead relies on the recognition of RNF8-generated K63-linked ubiquitin chains by the BRCA1-A complex through its subunit ABRAXAS (Sherker et al., 2021). Consistent with a contribution of the RNF168-dependent pathway, depletion of RNF168 using either an inducible shRNA system (Doil et al., 2009) or siRNA resulted in a partial, but not complete, reduction of BRCA1 and RAD51 recruitment to nucleolar caps (Figure 4C,D, Figure 4–figure supplement 1B–D). To test whether the RNF168-independent RAP80–ABRAXAS pathway also contributes to BRCA1 recruitment at nucleolar caps, we depleted RAP80 by siRNA. Strikingly, RAP80 depletion strongly impaired the formation of BRCA1 and RAD51 nucleolar caps in both U2OS and RPE1 cells following I-PpoI-induced rDNA damage (Figure 5A–D and Figure 5–figure supplement 1).

**Figure 3.**
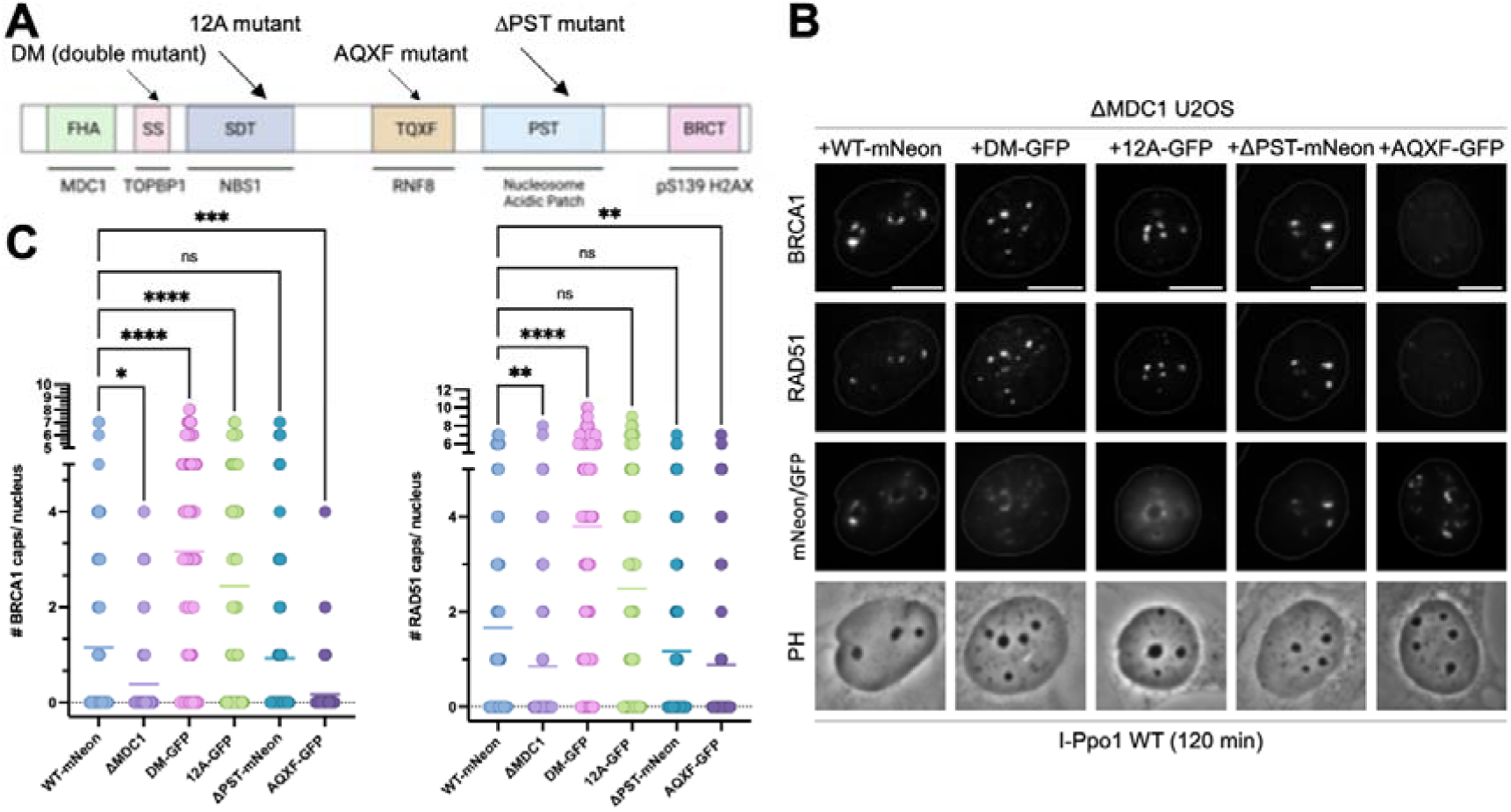
(A) A scheme of various MDC1 mutations introduced to ΔMDC1 U2OS cell line. (B) Localisation of BRCA1, RAD51, and mNeon/GFP-MDC1 after 2h of I-Ppo1 WT transfection. U2OS ΔMDC1 cells expressing wild type MDC1 (WT-mNeon), S168A/S196A MDC1 (DM-GFP), SDTADA MDC1 (12A-GFP), MDC1 lacking the entire PST repeat region (ΔPST-mNeon), and TQXFAQXF (AQXF-GFP). All scale bars = 10 μm. (C) Quantification of the experiment in (B) showing the number of BRCA1 (left panel) and RAD51 (right panel) nucleolar caps per cell (n= 100 cells). Graph represents a single replicate (experiment was performed in duplicates for RAD51 and in quadruplicate for BRCA1) and bars represent mean.

**Figure 4.**
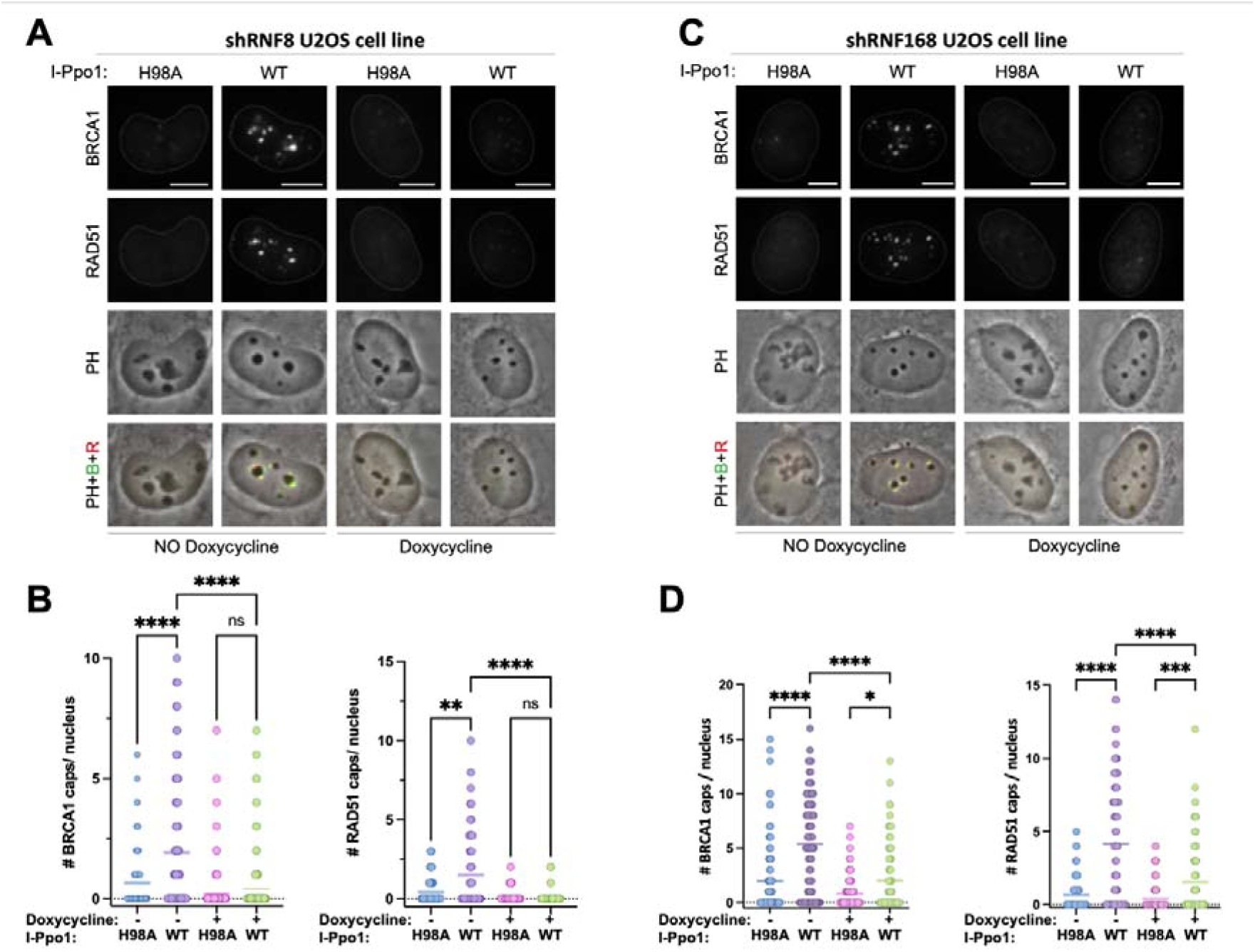
(A) Localisation of BRCA1 and RAD51 in doxycycline inducible RNF8 shRNA U2OS cell line treated with or without doxycycline after 2h of I-Ppo1 WT transfection. All scale bars= 10 μm. (B) Quantification of the experiment in (A) showing the number of BRCA1 (left panel) and RAD51 (right panel) nucleolar caps per cell (n= 100 cells). Graph represents a single replicate (experiment was performed in triplicates) and bars represent mean. (C) Localisation of BRCA1 and RAD51 in doxycycline inducible RNF168 shRNA U2OS cell line treated with or without doxycycline after 2h of I-Ppo1 WT transfection. All scale bars= 10 μm. (D) Quantification of the experiment in (C) showing the number of BRCA1 (left panel) and RAD51 (red panel) nucleolar caps per cell (n= 150 cells). Graph represents a single replicate (experiment was performed in triplicates) and bars represent mean.

**Figure 5.**
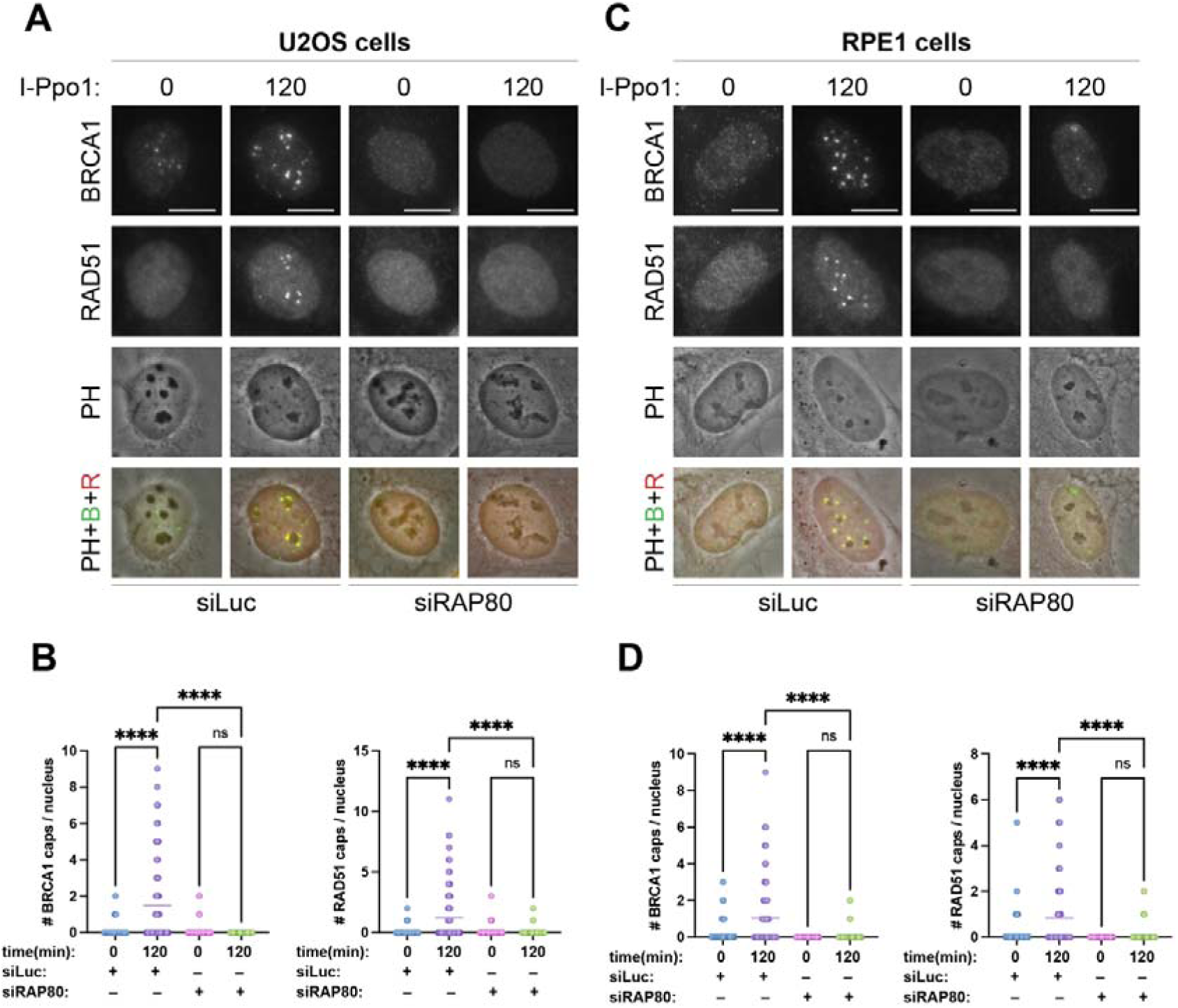
(A) Localisation of BRCA1 and RAD51 at baseline and after 2h of I-Ppo1 WT treatment in U2OScells transfected with siLuc or siRAP80. (B) Quantification of the experiment in (A) showing the number of BRCA1 (left panel) and RAD51 (right panel) nucleolar caps per cell (n= 138 cells). Graph represents one of two independent experiments and bars represent the mean. (C) Localisation of BRCA1 and RAD51 at baseline and after 2h of I-Ppo1 WT treatment in RPE1 cells transfected with siLuc or siRAP80. (D) Quantification of the experiment in (C) showing the number of BRCA1 (left panel) and RAD51 (red panel) nucleolar caps per cell (n= 105 cells). Graph represents one of two independent experiments and bars represent the mean. All scale bars = 10 µm

Together, these results indicate that BRCA1 recruitment to nucleolar caps depends on RNF8-mediated ubiquitylation and involves contributions from both the RNF168-dependent pathway and the RAP80–ABRAXAS complex, which together promote downstream RAD51 recruitment.

### The Treacle-NBS1-TOPBP1 complex mediates recruitment of the HDR machinery by acting upstream of γH2AX-MDC1-RNF8

So far, our data revealed that BRCA1 and RAD51 recruitment to nucleolar caps occur downstream of Treacle-NBS1-TOPBP1 and downstream of γH2AX-MDC1-RNF8. We therefore sought to determine if Treacle-NBS1-TOPBP1, and hence, rRNA transcriptional repression and nucleolar segregation, are required for H2AX phosphorylation and MDC1 recruitment. Indeed, downregulation of Treacle, TOPBP1 and NBS1 in U2OS by siRNA transfection resulted in almost complete abrogation of γH2AX and MDC1 signals within the nucleolar periphery 2h after transfection of cells with I-Ppo1 (Figure 6A,B). We previously showed that in response to targeted rDNA break induction, the Treacle-NBS1-TOPBP1 complex mediates ATM and ATR activation in the nucleoli (Mooser et al. 2020). Consistent with the idea that Treacle-NSB1-TOPBP1 acts upstream of γH2AX-MDC1-RNF8, we observed that pharmacological inhibition of ATM and ATR not only significantly reduced H2AX phosphorylation and MDC1 recruitment in response to rDNA breakage, but also strongly impaired recruitment of HDR factors to the nucleolar periphery (Figure 6–figure supplement 1A–E).

**Figure 6.**
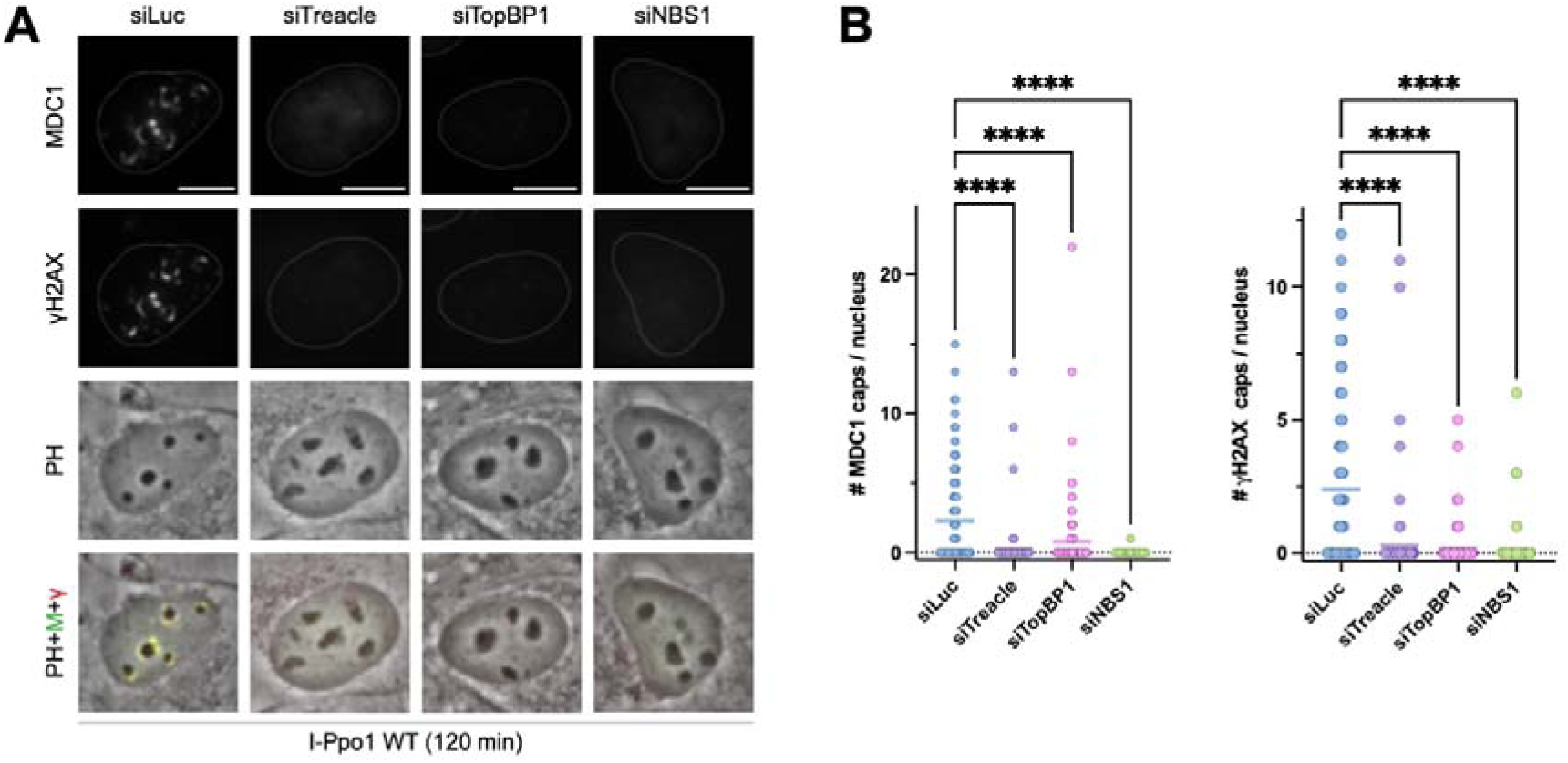
(A) Localisation of MDC1 and γH2AX in siLuc, siTreacle, siTopBP1, and siNBS1 transfected U2OS cells after 2h of I-Ppo1 WT treatment. All scale bars = 10 µm. (B) Quantification of the experiment in (A) showing the number of MDC1 (left panel) and H2AX (right panel) nucleolar caps per cell (n = 120 cells). Graphs represent a single replicate (experiment was performed in duplicates) and bars represent mean.

In summary, these data place Treacle-NBS1-TOPBP1 upstream, and recruitment of the HDR machinery downstream of γH2AX-MDC1-RNF8-dependent ubiquitylation pathways. To corroborate this model, we measured RAD51 recruitment to nucleolar caps in engineered BRCA1 (RPE1 ΔTP53 ΔBRCA1) and BRCA2 (DLD1 ΔBRCA2) knock-out cell lines and in two BRCA1 deficient breast cancer cell lines (MDA-MB-436 and SUM149PT). All of them show strong reduction of the RAD51 signal in the nucleolar periphery 2 h after transfection of the cells with I-Ppo1 (Figure 7A–D and Figure 7–figure supplement 1A–D). Moreover, at least within the first two hours after I-Ppo1 mRNA transfection, BRCA1 and RAD51 recruitment to nucleolar caps was cell cycle dependent, preferentially occurring in CyclinA positive S/G2-phase cells (Figure 7–figure supplement 1E,F). We also tested transcriptional repression upon rDNA break induction in RPE1 BRCA1 knock-out cells and detected no difference to the parental wild type cells, thus ruling out the possibility that the HDR machinery somehow regulates nucleolar segregation (Figure 7E,F).

**Figure 7.**
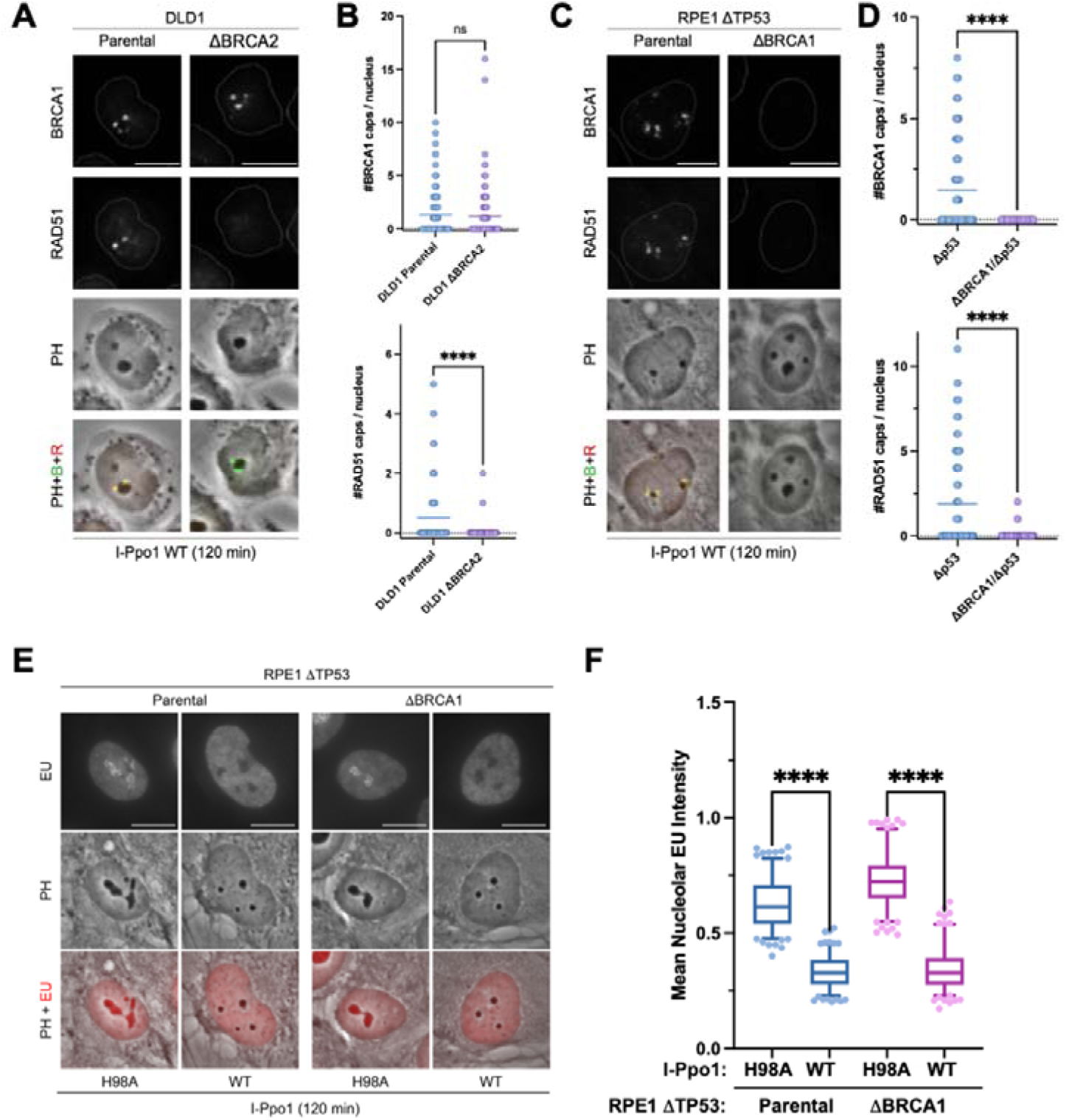
(A) Localisation of BRCA1 and RAD51 in parental and DLD1 ΔBRCA2 cells after 2h of I-Ppo1 WT treatment. All scale bars = 10 µm. (B) Quantification of the experiment in (A) showing the number of BRCA1 (upper panel) and RAD51 (lower panel) nucleolar caps per cell (n= 100 cells). Graph represents a single replicate (experiment was performed in duplicates) and bars represent mean. (C) Localisation of BRCA1 and RAD51 in RPE1 ΔTP53 and RPE1 ΔTP53 ΔBRCA1 cells after 2h of I-Ppo1 WT treatment. All scale bars = 10 µm. (D) Quantification of the experiment in (C) showing the number of BRCA1 (upper panel) and RAD51 (lower panel) nucleolar caps per cell (n = 100 cells). Graphs represent a single replicate (experiment was performed in duplicates) and bars represent mean. (E) Nucleolar EU incorporation in RPE1 ΔTP53 and RPE1 ΔTP53 ΔBRCA1 cells treated with I-Ppo1 WT or H98A for 2 hours. (F) Quantification of nucleolar EU incorporation after WT or H98A I-Ppo1 transfection in RPE1 ΔTP53 and RPE1 ΔTP53 ΔBRCA1 cells. Around 170 cells were considered per sample. Individual boxes represent the 25-75 percentile range with median and whiskers represent the 5-95 percentile range. Data points outside of this range are shown individually.

### MDC1 deficient cells are sensitive to rDNA damage and defective for DNA repair synthesis within nucleolar caps

So far, it was thought that MDC1 is not implicated in the nucleolar response to rDNA breakage (Larsen et al. 2014; Korsholm et al. 2020). However, we found that RPE1 ΔMDC1 cells are exquisitely sensitive to rDNA break induction, as sensitive as BRCA1 knock-out cells (Figure 8A,B). RPE1 H2AX^S139A^ knock-in cells are also sensitive to I-Ppo1 induced rDNA breaks, but to a lesser extent compared to ΔMDC1 and ΔBRCA1 cells (Figure 8A,B). Since we have shown that MDC1 is required for the recruitment of the HDR machinery into the nucleolar periphery, we were interested if MDC1 was also required for DNA repair at rDNA breaks. HDR is difficult to assess directly because no rDNA-specific HDR reporter system currently exists. However, HDR of DSBs involves DNA synthesis that can be measured by EdU incorporation. Yet, since we showed that BRCA1 and RAD51 are preferentially recruited to nucleolar caps in S/G2 phase cells (see above), it is difficult to distinguish EdU incorporation due to repair DNA synthesis from EdU incorporation due to replication DNA synthesis. To overcome this obstacle, we developed an experimental system based on *in situ* proximity ligation (PLA) using antibodies against γH2AX and incorporated EdU. This should allow the detection of EdU incorporation specifically at sites of DNA lesions around the nucleoli. Indeed, upon induction of rDNA breaks by I-Ppo1 wild type transfection, we observed a robust increase in the PLA signal around the nucleoli when both γH2AX and EdU antibodies were present in the reaction. This increase in the PLA signal upon rDNA break induction was significantly reduced in ΔMDC1 cells, which is consistent with impaired repair-associated DNA synthesis (Figure 8C,D). It should be noted though that we cannot rule out other effects that may lead to a decrease in the PLA signal in ΔMDC1 cells.

**Figure 8.**
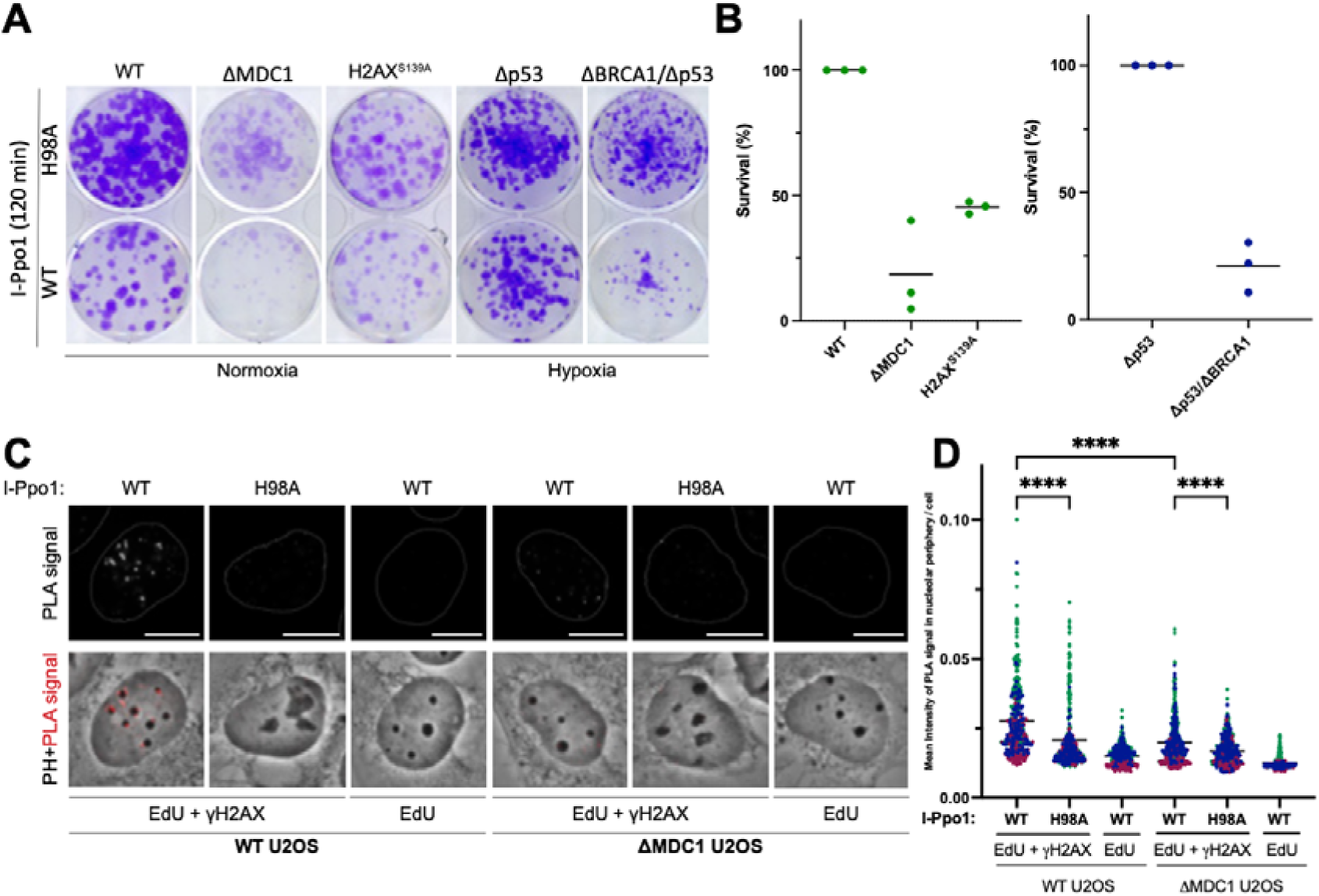
(A) Clonogenic survival assay of RPE1 WT, ΔMDC1 and H2AX^S139A^ cells as well PRE1 ΔTP53 (parental) and RPE1 ΔTP53 ΔBRCA1 cells after 2h of I-Ppo1 treatment. The RPE1 ΔTP53 and RPE1 ΔTP53 ΔBRCA1 cell lines were cultured under hypoxic conditions because BRCA1 deficiency confers sensitivity to atmospheric oxygen. (B) Clonogenic survival analysis of I-Ppo1 WT treated RPE1 WT, ΔMDC1 and H2AX^S139A^ or PRE1 ΔTP53 (Parental) and RPE1 ΔTP53 ΔBRCA1 cells. Each dot represents an independent replicate, black bars show means. (C) PLA assay showing the localization between EdU and γH2AX after H98A or WT I-Ppo1 transfection in U2OS or U2OS ΔMDC1 cells. All scale bars = 10 μm. (D) Quantification of the experiment in (C) showing the mean intensity of PLA signal within the nucleolar periphery U2OS or U2OS ΔMDC1 cells. Graph represents a pool of three independent experiments (n = 400 cells) and bars represent mean.

Together, these data indicate that DNA repair at rDNA breaks depends on γH2AX-MDC1 signaling, likely because these factors are required to establish RNF8-dependent chromatin ubiquitylation pathways that promote the recruitment of BRCA1 and subsequent accumulation of PALB2 and RAD51.

### Discussion

Here we propose that the recruitment of HRD factors to damaged rDNA repeats within nucleolar caps occurs downstream of two DDR adaptors that act sequentially: First, Treacle controls transcriptional repression and nucleolar segregation through phosphorylation-dependent interaction with NBS1 and TOPBP1, which promotes ATM and ATR activation. Once nucleolar segregation has occurred, MDC1 binds to phosphorylated H2AX within nucleolar caps and recruits HDR factors via RNF8-dependent chromatin ubiquitylation pathways involving both RNF168 and the RAP80–ABRAXAS complex (Figure 9). This dependency on γH2AX and MDC1 was unexpected because MDC1 has so far not been thought to be implicated in the nucleolar response to rDNA damage, even though it was shown to be recruited to nucleolar caps at late time points after targeted induction of rDNA breaks by CRISPR/Cas9 (Larsen et al. 2014; Korsholm et al. 2019). Instead, we and others have previously shown that NBS1 and TOPBP1, two proteins normally recruited to sites of DNA damage in an MDC1-dependent manner, were efficiently recruited to sites of nucleolar genotoxic stress in the absence of MDC1 by their direct and phosphorylation-dependent interaction with Treacle (Ciccia et al. 2014; Larsen et al. 2014; Mooser et al. 2020; Korsholm et al. 2019; Velichko et al. 2021). This previously led us to suggest that Treacle acted as the main adaptor for the nucleolar DDR, essentially taking the place of MDC1 (Mooser et al. 2020).

**Figure 9.**
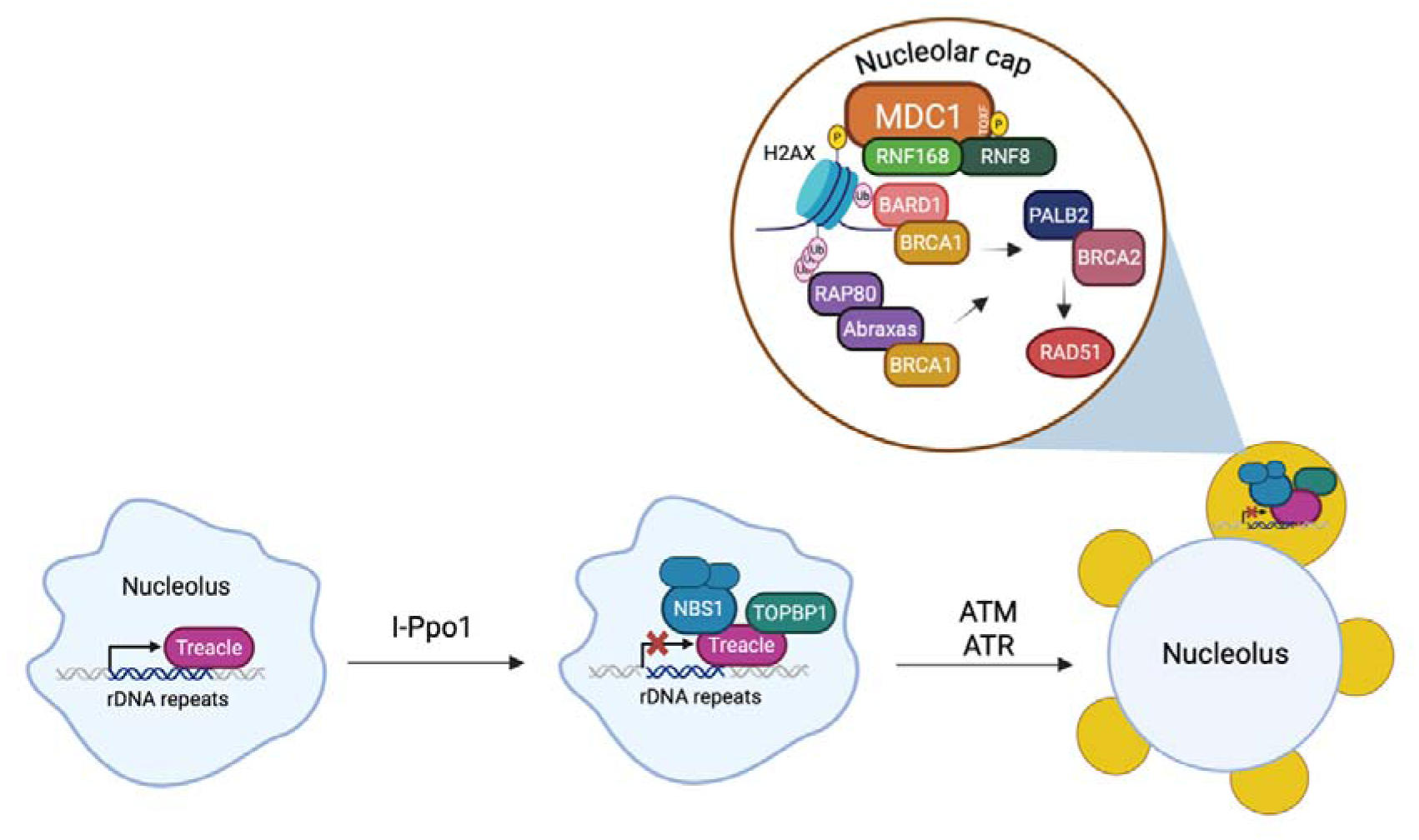
Proposed model of the nucleolar response to rDNA double-strand breaks. Before DNA damage, Treacle is constitutively associated with rDNA repeats (left). Following induction of rDNA double-strand breaks by I-PpoI, Treacle recruits the MRN complex and TOPBP1 through phosphorylation-dependent interactions, leading to ATM/ATR activation, transcriptional repression, and nucleolar segregation (middle). At nucleolar caps, H2AX is phosphorylated and recruits MDC1, which initiates RNF8-dependent chromatin ubiquitylation (right). RNF8 promotes BRCA1 recruitment through two parallel pathways: the RNF168–BARD1 pathway and the RAP80–ABRAXAS pathway. Together, these pathways promote the sequential recruitment of PALB2 and RAD51, enabling homologous recombination at the nucleolar periphery.

In this work, we would like to update this model. While MDC1 does indeed not seem to be required for the recruitment of NBS1 and TOPBP1 in the nucleoli, we show here that it is essential for the subsequent recruitment of DNA repair proteins in nucleolar caps, including key HDR factors.

It was recently shown that BRCA1 recruitment and RAD51 loading are controlled by two partially redundant pathways, both dependent on RNF8-mediated chromatin ubiquitylation (Sherker et al., 2021). One pathway is mediated by the RAP80–ABRAXAS complex, which recognises RNF8-generated K63-linked ubiquitin chains and promotes BRCA1 recruitment to damaged chromatin. The second pathway is mediated by the E3 ligase RNF168, which mono-ubiquitylates H2A on Lys15, thereby generating the H2AK15ub signal that is recognised by BARD1 to recruit BRCA1.

Our data indicate that both pathways contribute to HDR factor recruitment at nucleolar caps. While depletion of RNF168 resulted in a partial reduction of BRCA1 and RAD51 accumulation at nucleolar caps, depletion of RAP80 strongly impaired BRCA1 and RAD51 recruitment in two independent cell lines. Together, these findings support a model in which RNF8-dependent chromatin ubiquitylation promotes BRCA1 recruitment at nucleolar caps via both the RNF168-dependent and RAP80–ABRAXAS pathways, which together facilitate downstream RAD51 loading. This dual pathway mechanism closely mirrors BRCA1 recruitment at IR-induced DSBs (Sherker et al., 2021).

Our observation that RAD51 enrichment in nucleolar caps is dependent on MDC1 was surprising, given that γH2AX and MDC1 do not play a major role in RAD51 accumulation at sites of IR-induced breaks (Figure 2–figure supplement 1C), (Zhang et al. 2005; Scully and Xie 2013). Since both pathways for BRCA1 recruitment depend on RNF8-mediated chromatin ubiquitylation, it is possible that in the case of IR-induced breaks, RNF8 can also bind to damage sites independently of MDC1, which may be sufficient to recruit enough of the BRCA1-PALB2 axis to chromatin to efficiently load RAD51. Chromatin around rDNA breaks, on the other hand, which has to move first from inside of the nucleoli to the nucleolar periphery, may initially be less accessible for RNF8 and may thus require γH2AX-MDC1 for efficient priming by H1 poly-ubiquitylation.

Our observation that H2AX phosphorylation in response to targeted rDNA breaks is strictly dependent on Treacle-NBS1-TOPBP1 places this complex upstream of γH2AX. In contrast, there is currently no evidence that Treacle contributes to the repair of DSBs elsewhere in the genome, supporting the notion that its function is specialized for the nucleolar DNA damage response. Mechanistically, it is not yet clear why the Treacle-NBS1-TOPBP1 complex is required for H2AX phosphorylation. One possibility is that Treacle-NBS1-TOPBP1 is controlling ATM/ATR activation in the nucleoli. Consistent with this, we previously showed that upon rDNA breakage, both ATM and ATR are recruited in the nucleoli (Mooser et al. 2020). It is also possible that rDNA break mobilization must occur prior to H2AX phosphorylation. Limited H2AX phosphorylation was observed inside of the nucleoli, which may be explained by decreased nucleosome occupancy within the highly transcribed rDNA repeats (Korsholm et al. 2019; Marnef et al. 2019). In addition, H2AX phosphorylation was shown to spread beyond Treacle localization in nucleolar caps (Figure 2A,B), indicating that either rDNA is entering the perinucleolar heterchromatin compartment, or that additional chromatin regions become modified once nucleolar segregation has occurred. In any case, dependency of recruitment of HDR factors on Treacle-NBS1-TOPBP1, places HDR factor recruitment downstream of rDNA break mobilisation, which is tightly controlled by this complex. This may prevent premature RAD51 loading when NORs that contain broken rDNA repeats are still residing in the nucleoli and thus, are spatially juxtaposed to each other, which could result in recombination taking place between rDNA repeats that are located on different chromosomes. Consistent with this, we did not observe any RAD51 enrichment within the nucleoli at early time points after rDNA break induction, indicating that RAD51 loading only ensues once the NORs from different chromosomes have spatially segregated in the nucleolar periphery.

The spatio-temporal dynamics of DDR factor recruitment to sites of rDNA breaks is reminiscent of the dynamics of DDR factor recruitment to pericentric heterochromatic regions in mouse cells, which are concentrated in large structures within the nuclei termed chromocenters (Tsouroula et al. 2016). In both cases, HDR takes place at the periphery of the nuclear domains (chromocenters and nucleoli, respectively). There are, however, significant differences. Upon targeted induction of DSBs in pericentric heterochromatin, H2AX phosphorylation started at the periphery of the chromocenters and then spread to the core. In G1 phase cells, the γH2AX signal remained in the core, but in S and G2 phase cells it rapidly relocated to the periphery of the chromocenters (Tsuruoka et al. 2016). We did not observe similar cell cycle differences in γH2AX signal distribution in the nucleoli, indicating that regulation occurs primarily via temporal rather than spatial dynamics of H2AX phosphorylation.

It was proposed that DSBs in rDNA repeats and centromeric heterochromatin can be repaired by HDR in G1 (Yilmaz et al. 2021; van Sluis and McStay 2015). Our analysis revealed that BRCA1 recruitment and RAD51 loading preferentially took place in S/G2 phase cells, at least within the first 2 hours after I-Ppo1 mRNA transfection. This agrees with our previous study where we showed that DNA end resection of broken rDNA repeats also preferentially takes place in S/G2 cells (Mooser et al. 2020). We propose that cell cycle independent HDR only takes place at later time points, at persistent rDNA breaks, which may be resected also in G1-phase cells. Since we also observed recruitment of 53BP1 to nucleolar caps and hyper-resection in cells depleted of 53BP1 (data not shown), we surmise that cell cycle regulation of HDR at DSBs within rDNA repeats may be analogous to that observed at IR-induced breaks. Further analysis such as cell cycle synchronization or use of phase-specific markers will be necessary to confirm this idea.

Together, our findings establish a mechanistic framework in which nucleolar reorganization and RNF8-dependent chromatin ubiquitylation are tightly coordinated to ensure spatially controlled recruitment of HDR factors to rDNA breaks. This coupling of DNA repair to subnuclear compartmentalization may represent a general strategy to safeguard genome stability within repetitive DNA regions and to limit aberrant recombination events. Although Treacle is encoded by TCOF1, the gene mutated in Treacher Collins syndrome, the developmental defects associated with this disorder are generally attributed to impaired ribosome biogenesis and nucleolar homeostasis rather than defective nucleolar DNA damage signaling. Whether the DNA damage response function of Treacle contributes to disease pathology remains an interesting question for future studies.

## Acknowledgements

We thank Steve Jackson, Jiri Lukas, Niels Mailand, Brian McStay and Lorenza Penegno for providing valuable reagents. Imaging was performed with equipment maintained by the Center for Microscopy and Image Analysis, University of Zürich. Cell sorting was carried out by the Flow Cytometry Core Facilities at the University of Zürich. The Stucki lab is supported by two project grants from the Swiss National Science Foundation (31003A_163141 and 310030_189141) and by the Kanton of Zü□rich.

## Author Contributions

Andrea Hänel, Conceptualization, Data curaction, Formal analysis, Investigation, Supervision, Methodology, Writing – original draft; Johannes Leyrer, Data curation, Formal analysis, Supervision, Investigation, Methodology; Polyxenia Stutz, Data curation, Formal analysis, Methodology; Manuel Stucki, Conceptualization, Supervision, Validation, Investigation, Writing – original draft, Project administration, Writing – review and editing.

## Materials & Methods

### Cell culture and drug treatment

All cell lines were grown in a sterile cell culture environment and routinely tested for mycoplasma contamination. U2OS (human osteosarcoma cell line), RPE1 (immortalized human retinal pigment epithelial 1 cell line), HeLa (cervical carcinoma cell line), and 293T (transformed human embryonic kidney cell line) cells were cultured in Dulbecco’s modified Eagle medium (DMEM) supplemented with 10% fetal bovine serum (FBS), 2mM L-glutamine, and penicillin-streptomycin (PenStrep) antibiotics under standard cell culture conditions in a CO_2_ incubator (37 °C, 5% CO_2_). p53 and BRCA1/p53 RPE-1 cells were cultured using hypoxia chamber (4.96 % mol CO_2_, 3.15 % mol O_2_, 91.89% mol N_2_) and standard DMEM + 10% FBS and 5% PenStrep media. Parental and ΔBRCA2 DLD1 (colorectal adenocarcinoma cell line), and MDA-MB436 (breast adenocarcinoma cell line) cells were cultured in RPMI (Roswell Park Memorial Institute)-1640 medium supplemented with L-glutamine, 10% FBS and 5% PenStrep medium. SUM149PT (invasive breast cancer cell line) cells were cultured in Ham’s F12 medium supplemented with L-glutamine, 10% FBS and 5% PenStrep. Stably expressing doxycycline-inducible RNF8 shRNA or RNF168 shRNF168 U2OS cells were cultured in the presence of 1µg/ml Puromycin (Invivogen) and 5 µg/ml Blasticidin (Thermo Fisher Scientific) in DMEM medium + 10% FBS (without tetracycline) + 5% PenStrep. The expression of shRNF8 or shRNF168 was induced by 1 µg/ml doxycycline (Clontech) for 48 h or 72 h, respectively. U2OS cell lines stably transfected with MDC1 WT-mNeon expression constructs were cultured in the presence of 25 µg/ml Hygromycin (Life Technology). MDC1 PST-mNeon U2OS cells were cultured in the presence of 1 µg/ml Puromycin (Invivogen). MDC1 12A-GFP, MDC1 DM-GFP, and AQXF-GFP U2OS cell lines were cultured in the presence of 400 µg/ml G418 (Sigma). U2OS cell lines stably transfected with Treacle-GFP and tetracycline repressor expression constructs were cultured in the presence of 200 μg/ml Zeocin (Life Technology) and 10 μg/ml Blasticidin (Thermo Fisher Scientific).

For pulsed EU incorporation, cells were incubated for 30 min in medium without antibiotics containing 1 mM EU. The Click-iT EU Alexa Fluor 594 Imaging Kit (Thermo Fisher Scientific) was used for EU detection. For pulsed EdU incorporation, cells were incubated for 1 h in medium without antibiotics containing 20 µM EdU. ATM inhibitor KU- 55933 (Selleckchem) and ATR inhibitor VE-821 (Selleckchem) were used in the final concentration of 5μM for 4h. X-ray irradiation of cells was performed using Xylon.SMART 160E-1.5 machine employing 2 Gy with the recovery time of 2h.

### Stable cells lines and cloning

The generation of stable cell lines used in the project is described in detail in corresponding references: Treacle-GFP U2OS, NBS1 U2OS, and NBS1-mNeon U2OS (Mooser et al. 2020). ΔMDC1 U2OS, MDC1-mNeon U2OS, MDC1-12A-GFP U2OS, MDC1-AQXF-GFP U2OS, and MDC1-DM-GFP U2OS (Leimbacher et al. 2019) PST-MDC1-mNeon was generated in our laboratory as described in using Quick Change II XL- Site directed mutagenesis kit (Agilent) and following primers; forward: p-TTT ACC CCC ACA GAC CAG, Reverse: p-AGC AGA GGT AGC TGG AAA. ΔMDC1 HeLa was generated in our laboratory using a sense and an antisense guide RNA targeting the FHA domain of MDC1 with minimal off-target impact, which were introduced in the All-In-One (AIO) CRISPR-Cas9D10A-mCherry plasmid (Addgene) described in (Chiang et al. 2016). In detail, the sense gRNA was cloned between the BsbI sites before the first gRNA scaffold and the antisense gRNA was cloned was cloned between BsaI sites before the second gRNA scaffold resulting in the AIO CRISPR-Cas910A-mCherry-MDC1-gRNA plasmid, which was verified by Sanger sequencing. This plasmid was used to transfect HeLa cells together with jetOPTIMUS® DNA transfecting reagent. After 48 h, mCherry-positive single cells were sorted into 96-well plates and individual wells containing single colonies were identified using transluminescence microscopy. Colonies were progressively expanded and the absence of MDC1 was verified by both, western blotting and Sanger sequencing. MDC1 RPE-1 and H2AX^S139A^ RPE-1, a kind gift of Steve Jackson, were previously described (Chiang et al. 2016).

### RNA and DNA transfection

The control siRNA (siLuc), siRNA targeting Treacle (siTreacle), TopBP1 (siTopBP1), NBS1 (siNBS1) and BRCA1 (siBRCA1) were obtained from Microsynth AG. The siRNA pools targeting RNF168 and RAP80 were optained from Dharmacon Reagents (SMARTPool ON-TARGETplus siRNA). The sequences of the siRNAs are the following:

siLuc: UGGUUUACAUGUCGACUAA-dTdT

siTreacle: CCACCAUGGGUUGGAACUAAAUU-dTdT

siTopBP1: ACAAAUACAUGGCUGGUUA-dTdT

siNBS1: GGAGGAAGAUGUCAAUGUUTT

siBRCA1: GGAACCUGUCUCCACAAAFTT

For siRNA transfection, cells were grown in a six-well plate 24 h prior to transfection. A total of 60-70% confluent cells were transfected using 20 nM siRNA, or 30 nM siRNF168 respectively, and Lipofectamine RNAiMAX^TM^ (Invitrogen) according to the manufacturer’s instructions. Cells were incubated with siRNA-lipid complexes for 48 h before being either seeded on cover slips (immunofluorescence) or expanded (Western blot) and harvested 72 h after siRNA transfection. I-Ppo1 mRNA transfection was done using Lipofectamine MessengerMaxTM Reagent (Invitrogen) according to the manufacturer’s protocol. For the generation of the I-Ppo1 endonuclease, we used the vectors pIRES I-Ppo1 wild type and H98A (a kind gift from Brian McStay) (van Sluis and McStay 2015). Plasmids were linearized with NotI and transcribed *in vitro* using the MEGAscript T7 kit (Ambion) according to the manufacturer’s instructions. The I-Ppo1 mRNA was then polyadenylated using the Poly(A) tailing kit (Ambin) according to the manufacturer’s instructions. For I-Ppo1 transfection, cells were grown on covers slips or in six-well plate 24 h prior to transfection. Cells (at around 80% confluency) were transfected using 500 ng of I-Ppo1 mRNA per cover slip (24-well plate) or 1500 ng of I-Ppo1 mRNA per 6-well plate and Lipofectamine MessengerMax^TM^ Reagent (Invitrogen) according to the manufacturer’s instruction. Plasmid DNA (pcDNA4-TO-strep-HA-GFP-Treacle) for immunoprecipitation experiment was transfected using jetOPTIMUS® DNA Transfection Regent in 6-well plates following the standard protocol provided by the manufacturer. Cells were then incubated for 24 h before proceeding further.

### SDS-PAGE and western blot

To prepare cell extracts for SDS-PAGE, cells were washed in PBS and scraped off the plate using standard SDS buffer (120mM Tris pH 6.8, 20% glycerol, 4% SDS). Cell extracts were sonicated for 2 x 30 s, and the final protein concentration was measured using NanoDrop. Protein samples (50 μg) were mixed with 2x SDS loading dye (100 mM DL-Dithiothreitol, 4% SDS, 20% glycerol, 0.2% bromophenol blue, 100 mM Tris-Cl (6.8)) and heated for 5 min at 95 °C before loading on a gel. SDS-PAGE was performed using 4–20% Mini-PROTEAN TGX Stain-free Precast Gels (Bio-Rad). As a reference for molecular weights, Prestained Protein Ladder 245 kDa (Geneaid) was loaded on a gel. Western blotting was performed using the Trans-Blot® TurboTM Transfer system (Bio-Rad) on Mini Format 0.2 μm nitrocellulose membranes (Bio-Rad). For detection of BRCA1 protein, 0.2 μm PVDF membrane (Bio-Rad) was used, and membrane blocking was performed with 5% milk in TBS-Tween for at least 2 hours. All nitrocellulose membranes were blocked using EveryBlot Blocking Buffer (Bio-Rad) for at least 10 min except for membrane for the RNF8 and RNF168 detection (5% milk in TBS-Tween was used). The following antibodies were used in the indicated dilutions: BRCA1 (Mouse, Santa Cruz, SC6954, 1:500), BRCA2 (Mouse, Merck, OP95-100UG, 1:500), Treacle (Rabbit, Atlas Antibodies, HPA038237, 1:500), TopBP1 (Rabbit, Abcam, ab2402, 1:500), NBS1 (Rabbit, Merck, JBW301, 1:1000), MDC1 (Rabbit, Abcam, ab11171, 1:5000), Chk1 (Mouse, Santa Cruz, sc-8408, 1:1000), pS317-Chk1 (Rabbit, Cell Signaling, 2665, 1:500), Chk2 (Rabbit, Cell Signaling-Bioconcept, 2662S, 1:1000), pT68-Chk2 (Rabbit, Cell Signaling, 2661, 1:500), RNF8 (kind gift from Prof. Penengo), RNF168 (Rabbit, Thermo Fisher Scientific, PA5-65427, 1:1000), RAP80 (Rabbit, Bethyl, A300-763A, 1:5000), Tubulin hFAB^TM^ Rhodamine, (Biorad, 12004166, 1:1000), HA-HRP (Chicken, Abcam, ab1190, 1:500). Blots were incubated with primary antibodies overnight at 4 °C and washed 3 × 10 min with TBS-Tween before incubation for 1 h with secondary antibodies at room temperature. Following secondary antibodies were used: HRP- conjugated anti-mouse (GE Healthcare, NA934, 1:2500) or HRP-conjugated anti- rabbit (GE Healthcare, NA934, 1:2500). Blots were developed with SuperSignal^TM^ West Femto Maximum Sensitivity Substrate (Thermo Fisher Scientific) and image acquisition was done on a ChemiDoc MP Imaging System (Bio-Rad).

### Immunoprecipitation

To prepare cell extracts for immunoprecipitation, cells were washed 1 x with PBS and harvested using cell scrapers. After centrifugation at 2300 g for 3 min at 4 °C, supernatant was removed, and cells were resuspended in 100-200 μl (depending on the number of cells) of IP buffer (50 mM Tris-HCl pH 7.6, 100 mM NaCl, 1 mM MgCl_2_, 10% glycerol, 5 mM NaF, 0.2% NP-40, 2 mM EDTA), supplemented with complete EDTA-free protease inhibitor cocktail (Roche) and 25 U/ml Benzonase (Sigma). Cell lysates were incubated on ice for 30 min and centrifuged at 13 000 rpm for 15 min at 4 °C. For each IP reaction, 30 μl of monoclonal anti-HA agarose beads (Sigma, A2095) were added to 1 mg of the soluble cell extract and samples were incubated overnight with end-over-end rotation at 4 °C. Next day, cells were washed three times with IP buffer and resuspended in 2x SDS loading dye (100 mM DL-Dithiothreitol, 4% SDS, 20% glycerol, 0.2% bromophenol blue, 100 mM Tris-Cl (pH 6.8)).

### Clonogenic survival assay

RPE-1 cells were counted using automated cell counter Countess II (Thermo Fisher Scientific) and plated in 24-well plates at a density of 8*10^3^ cells per well. The day after seeding, cells were transfected with I-Ppo1 WT or H98A mRNA for 2 h prior to medium exchange. To ensure the same conditions, cells were counted and plated in triplicates in 6-well plates at a density 1000 cells per well. Colonies were grown for 12 days, with medium change every 3 days. At the assay endpoint, colonies were washed with PBS and fixed with 4% formalin for 12 min before staining with 0.5% crystal violet in 10% EtOH solution (Honeywell) for 10 min at RT. After rigorous washing with PBS, plates were left to dry overnight. Images of the colonies were acquired using flatbed scanner (Epson perfection V800). Quantification was performed using the Colony-Area plug-in of Fiji40.

### Immunofluorescence

Cells were grown on glass coverslips and washed with PBS before fixation with 4% formalin for 12 min at RT. Formalin was discarded and cells were washed 3 x 5 min with PBS at RT and subsequently permeabilized for 5 min in PBS containing 0.3% Triton X. After another round of washing with PBS, cells were incubated with blocking buffer (10% FBS in PBS) for at least 1 h at RT. Primary antibody incubations were performed at 4°C overnights. The next day, cover slips were washed 3 x 5 min with PBS and secondary antibody staining was performed for 1h at RT in the dark. After washing with PBS (at least 3 x 10 min), coverslips were mounted on glass microscopy slides with VECTASHIELD® PLUS Antifade Mounting Medium containing 0.5 μg/mL 4’,6-diamidino-2-phenylindole dihydrochloride (DAPI; Vector Laboratories). The following antibodies were used at the indicated dilutions: CtIP (Rabbit, LubioScience, A300-488A-T, 1:100), BRCA1 (Mouse, Santa Cruz Biotechnology, sc-6454, 1:100), RAD51 (Rabbit, Santa Cruz Biotechnology, sc-8349, 1:500), PALB2 (Rabbit, Invitrogen, PA5-66053, 1:600), MDC1 (Rabbit, Abcam, ab11171, 1:500), γH2AX (Mouse, Millipore, 05-636, 1:500), Treacle (Rabbit, Sigma Life Science, HPA038237, 1:100), Cyclin A (Rabbit, Santa Cruz Biotechnology, sc-751, 1:100), Cyclin A (Mouse, BD Bioscience, 611269, 1:100), GFP-Booster Alexa 488 (Chromotech, gb2AF488-10, 1:500). The following secondary antibodies were used: Alexa Fluor 488 Goat Anti-Mouse (Life Technologies, A11029, 1:1000), Alexa Fluor 568 Goat Anti-Mouse (Life Technologies A11031, 1:1000), Alexa Fluor 647 Goat Anti-Mouse (Life Technologies A21235, 1:1000), Alexa Fluor 488 Goat Anti-Rabbit (Life Technologies A11034, 1:1000) and Alexa Fluor 568 Goat Anti-Rabbit (Life Technologies, A11036, 1:500).

### PLA assay

U2OS WT and MDC1 were seeded at 80.10^3^ on cover slips in medium without antibiotics and I-Ppo1 WT or H98A was added the next day. After 1h, EdU was added on cover slips (20 µM) and cells were fixed with formalin (12 min at RT) after another hour. Cells were washed 3 x 5 min with PBS and subsequently permeabilized for 5 min in PBS containing 0.3% Triton X at RT. After another round of washing with PBS, cells were incubated with 1x blocking solution (Sigma Duolink PLA kit) in a pre-heated humidity chamber for 1.5 h at 37°C. Blocking solution was removed, and cells were incubated for 30 min in a EdU Click-iT solution containing biotin azide according to a manufacture’s protocol. Next, cells were incubated with primary antibodies; Biotin (Rabbit, Bethyl LubioScience, A150-109A, 1:1250) and γH2AX (Mouse, Millipore, 05-636, 1:500) in a humidity chamber over night at 4°. The next day, cells were washed 2 x 5 min with wash buffer A at RT and subsequently incubated with MINUS or PLUS PLA probes (Sigma Duolink PLA kit) for 1 h at 37° in a pre-heated humidity chamber. Cells were then washed 2 x 5 min with wash buffer A and incubated with a ligation solution (1x ligation buffer and ligase enzyme from the Sigma Duolink ligation kit) for 30 min at 37°C. Cells were then washed 2 x 5 min with wash buffer A and incubated for 100 min at 37° with an amplification solution (1x amplification buffer and polymerase enzyme from the Sigma Duolink amplification kit). Subsequently, cells were washed 2 x 10 min with wash B, 1 x 5 min with ddH_2_O and coverslips were mounted on glass microscopy slides with VECTASHIELD® PLUS Antifade Mounting Medium containing 0.5 μg/mL DAPI (Vector Laboratories).

### Widefield and super-resolution microscopy

Widefield image acquisition was done on a Leica DMI6000B inverted fluorescence microscope, equipped with Leica K5 sCMOS fluorescence camera (16-bit, 2048 × 2048-pixel, 4.2 MP) and Las X software version 3.7.2.22383. HCX Plan Apochromat 63X/1.40 and Plan Apochromat 100X/1.40 phase contrast oil immersion objectives were used for image acquisition. For quadruplet-wavelength emission detection, the combination of DAPI with GFP (m-Neon) or Alexa Fluor 488, Alexa Fluor 568, and Alexa Fluor 647 was used. For structured-illumination high-resolution confocal microscopy cells were grown on high precision glass coverslips #1.5H, 0.17 mm thick (Assistent) and processed for immunofluorescence as described above. Cells were then stained with 1 μg/mL DAPI (AppliChem, A1001) diluted in PBS for 10 min at room temperature in the dark. DAPI was discarded and cells were further washed with deionized water to completely remove PBS. Coverslips were mounted on microscopy glass slides without frosted edges (R. Langenbrick GmbH, Dimensions L 76 × W 26 mm) using VECTASHIELD® HardSetTM Antifade Mounting Medium (Vector Laboratories). Structured illumination confocal imaging was carried out at the ScopeM imaging facility of ETH Zurich, using DeltaVision OMX microscope equipped with DeltaVision OMX (3.30.4378.0) software, PlanApoN 60x /1.42NA oil PSF objective, and sCMOS OMX V4 (15bit range) cameras. DAPI was detected using 405 nm diode laser (Vortran Stradus^TM^) and 420-451 nm emission filters, GFP and Alexa 488 signal was detected with the 488 nm vertical external cavity surface emitting laser (VECSEL) and 504-553 nm emission filters, and Alexa 568 was detected using 568 nm emitting laser (VESCEL) and 590-627 nm emission filters. Z-stacks of entire cells were acquired using optimal step size. The reconstruction of OMX images was done using softWorx software 6.0 (5.9.9). The approximate lateral resolution was 120-160 nm, and the axial 280-350 nm. For optimal representation of confocal images in figures, maximum intensity projection was calculated using Fiji. The displayed intensity of images was adjusted for optimal demonstration of structures of interest using Adobe Photoshop.

### Image quantification and processing

Nucleolar caps quantification after rDNA damage was done using Cell Profiler 4.0.6. First, nuclei segmentation was performed by the intensity-based primary object detection module using DAPI signal. Next, nucleoli were detected by enhancing dark holes in the phase contrast images followed by detection of primary objects based on their intensities. Area, in which nucleolar caps were detected, was created by expanding the identified nucleoli and by shrinking the identified nucleoli. The two objects were masked and a torus around a nucleolus was created. Nucleolar caps were segmented by intensity-based primary object detection module using the fluorescence signal of the protein of interest. Detected objects were masked with the designated areas around individual nucleoli and the object area, shape, and intensity were measured. Based on the measurements, objects were filtered to exclude background and random foci. Finally, objects were related, and the area, shape, and intensity of nucleolar caps were quantified. For quantification of the EU signal in WT RPE-1 cells and p53 and BRCA1/p53 RPE-1-hTERT cells, a simple pipeline in Cell profiler 4.0.6, measuring the intensity of EU signal inside of nuclei, was created. For quantification of the PLA experiment, Cell Profiler was employed, and the intensity of PLA signal was measured in the designated areas around individual nucleoli, which was created as described above. For experiment, where IR-induced foci were quantified, intensity-based identification of primary object within a Dapi positive region was used. Finally, Cell Profiler was used to measure nucleolar caps or IR-induced foci in cells, which were grouped based on the intensity of Cyclin A signal. The quantification of BRCA1 and RAD51 positive foci in NBS1 and WT U2OS cells as well as PALB2 positive foci in siBRCA1 U2OS treated cells was performed manually counting cells positive for BRCA1/RAD51/PALB2 nucleolar caps. The 3D reconstruction of SIM images was performed using the 3D rendering feature of the Imaris software package. The processing of all microscopy images was done using Adobe Photoshop and the final figures were created in Adobe InDesign.

### Statistical Analysis

All graphs were generated using GraphPad Prism 9 or 10. The appropriate statistical test was chosen as follows: Unpaired non-normal distributed data of two groups were tested with two-tailed Mann–Whitney test. Unpaired normal distributed data of two groups were tested with Sidak’s test. Three and more groups were analysed by one-way ANOVA. In case of non-normal distribution, Kruskal-Walli’s test for multiple comparison was used. All statistical tests were run with alpha error level 0.05 and significance was considered using p value (p < 0.0001 means ****). Unless otherwise indicated, horizontal bars represent the mean values and error bars the standard deviations. In box plot graphs, boxes represent the 25–75 percentile range with median, and whiskers represent the 9–95 percentile range. Data points outside this range are shown individually.

