## Supplementary material for "Sequential adaptor function of Treacle and MDC1 couples nucleolar reorganization to RNF8-dependent recruitment of HDR factors": Figure supplements

**Figure 1–figure supplement 1**

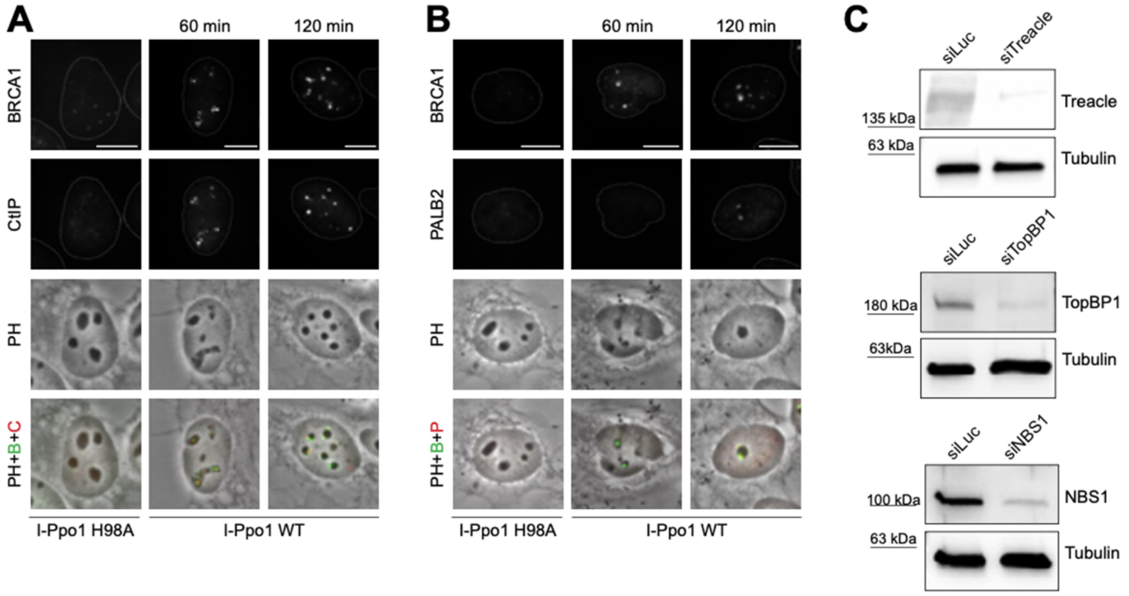


(A) Lo calisation of CtIP and BRCA1 after 1h or 2h of I-Ppo1 WT transfection or after 2h of I-Ppo1 H98A transfection. (B) Localisation of PALB2 and BRCA1 after 1h or 2h of I-Ppo1 WT transfection or after 2h of I-Ppo1 H98A transfection. All scale bars = 10 $\mu$m. (C) Western blotting of extracts from U2OS cells treated with siLuc (control), siTreacle, siTopBP1, and siNBS1.

Figure 1–figure supplement 2

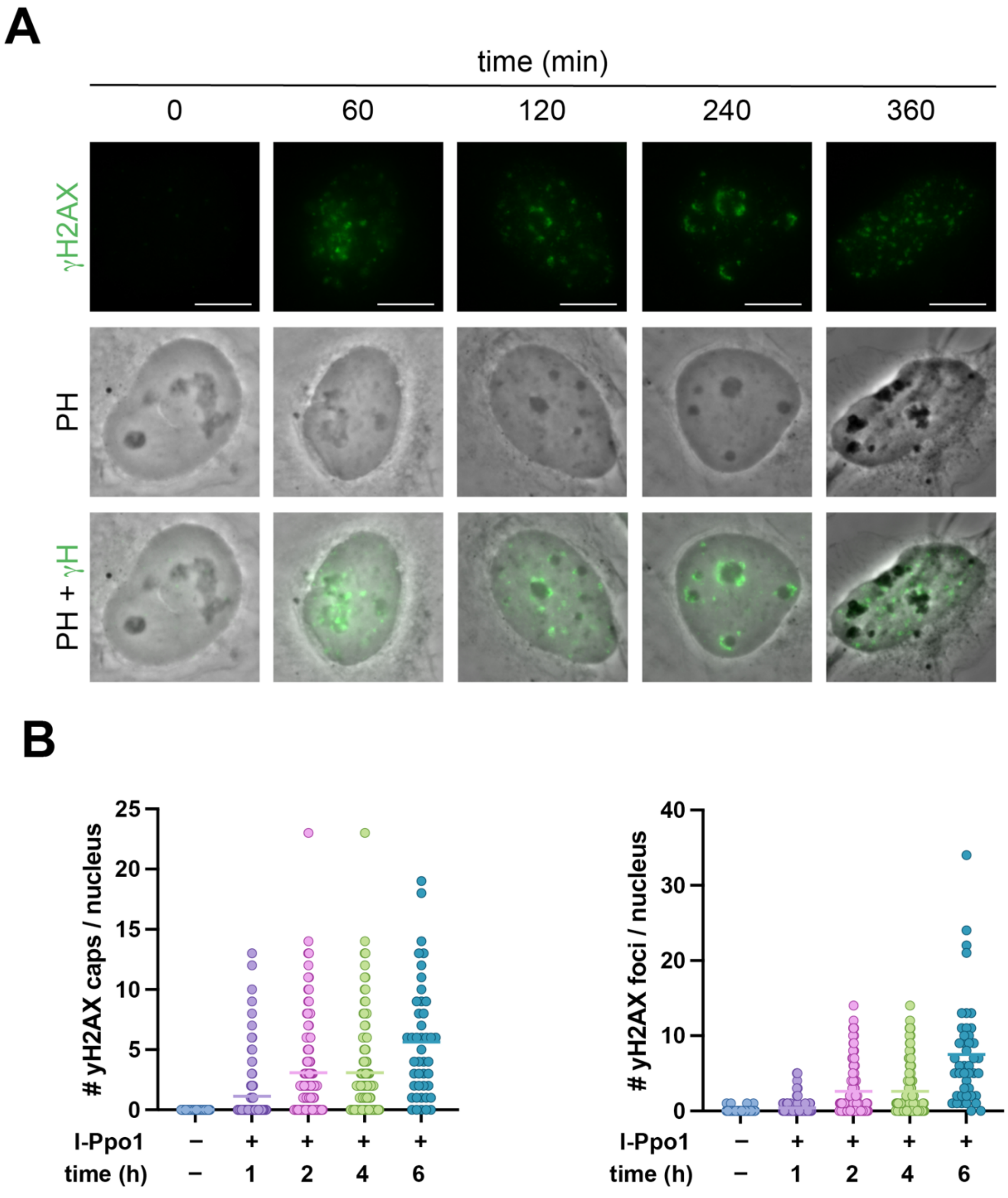


(A) Localization γH2AX foci at baseline and at various time points after transfection with I-Ppo1 WT. (B) Quantification of the experiment in (A). Graph represents one of two independent experiments and bars represent the mean. Scale bars = 10 $\mu$m

**Figure 1–figure supplement 3**

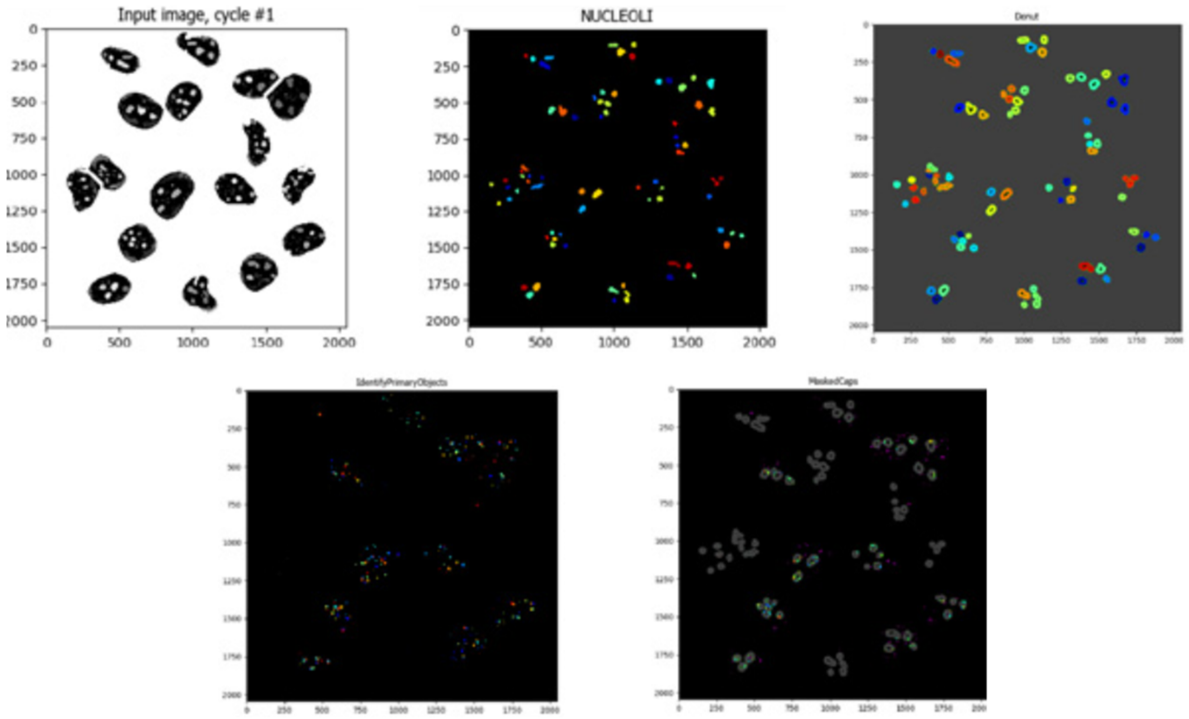


Cell Profiler pipeline for quantification of caps in the nucleolar periphery after rDNA DSBs induction.

**Figure 2–figure supplement 1**

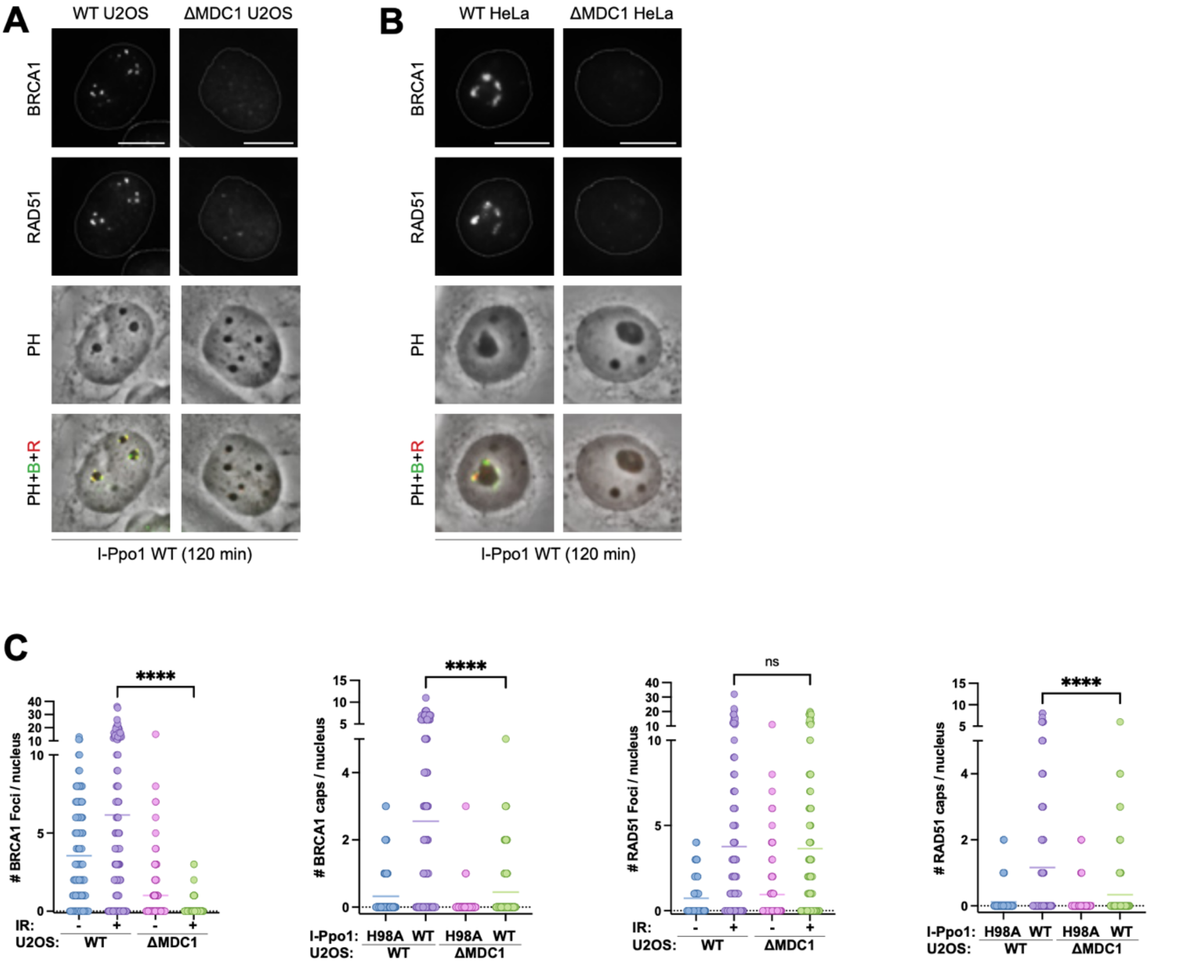


(A) Localization of BRCA1 and RAD51 in WT and ΔMDC1 U2OS cells after 2h of I-Ppo1 WT transfection. (B) Localization of BRCA1 and RAD51 in WT and ΔMDC1 HeLa cells after 2h of I-Ppo1 WT transfection. All scale bars = 10 $\mu$m (C) Quantification showing the number of BRCA1 foci (first panel) or the number of BRCA1 nucleolar caps (second panel) per nucleus (n= 100), and the number of RAD51 foci (third panel) or the number of RAD51 nucleolar caps (fourth panel) per nucleus (n= 100) in WT or ΔMDC1 U2OS cells treated with or without irradiation, or with I-Ppo1 WT or H98A, respectively. Graphs represent a single replicate, and bars represent mean.

**Figure 3–figure supplement 1**

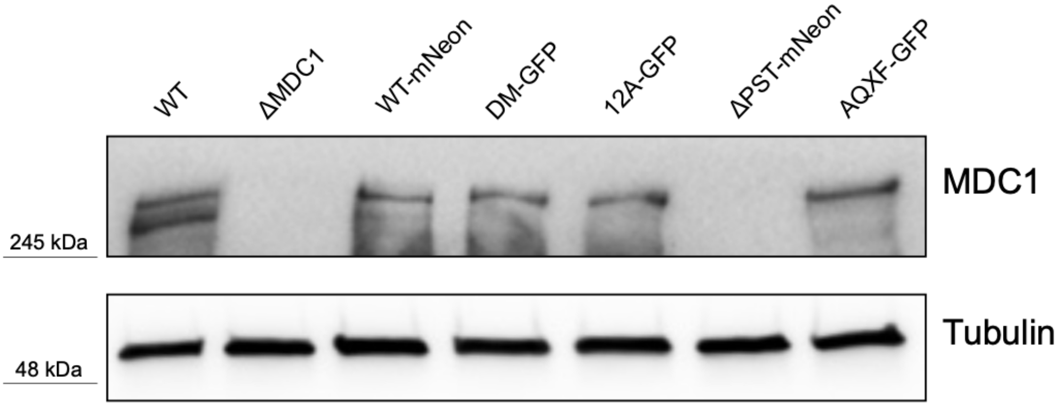


(A) Western blotting of extracts from WT, WT-mNeon, ΔMDC1, DM-GFP, 12A-GFP, ΔPST-mNeon, and AQXF-GFP U2OS cells. Note that there is no signal in ΔPST-mNeon becaue the antibody was raised against a fragment from the PST repeat region

**Figure 4–figure supplement 1**

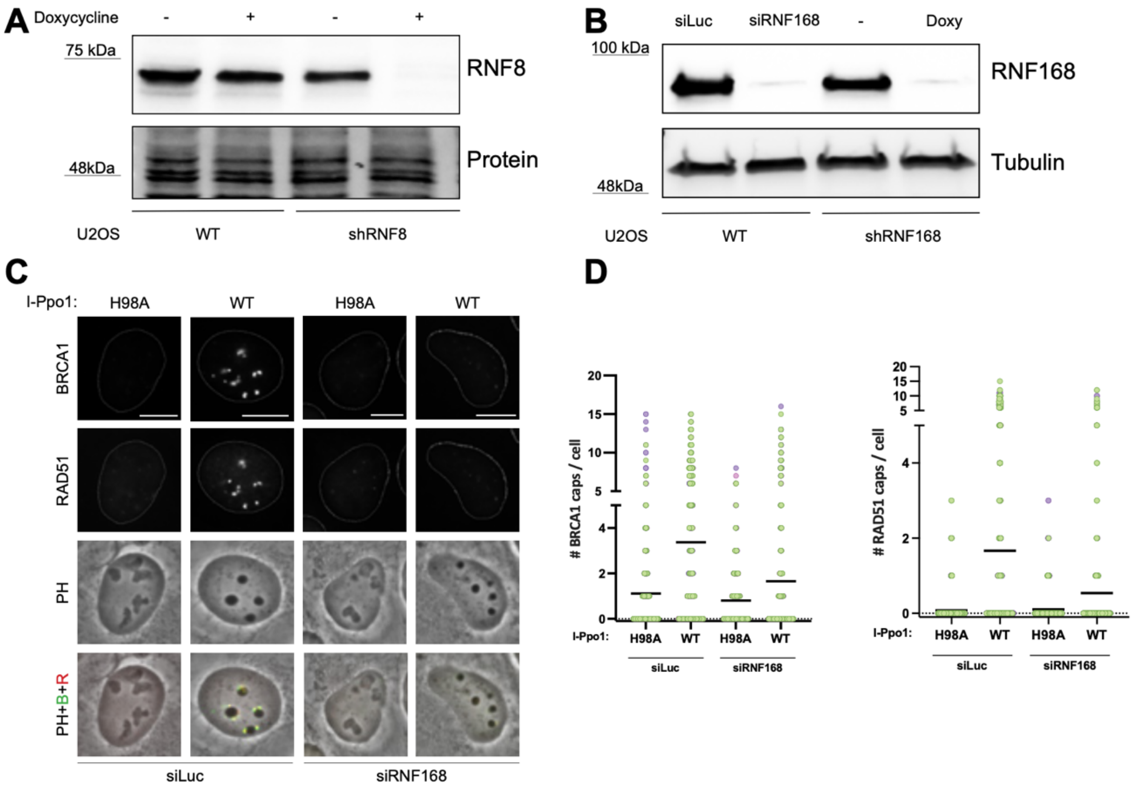


(A) Western blotting of extracts from WT and doxycycline inducible RNF8 shRNA U2OS cells treated with or without doxycycline. (B) Western blotting of extracts from siLuc (control) or siRNF168 treated U2OS WT cells and extracts from doxycycline inducible RNF168 shRNA U2OS cells treated without or with doxycycline. (C) Localisation of BRCA1 and RAD51 in siLuc and siRNF168 transfected U2OS cells after 2h of I-Ppo1 WT or H98A treatment. All scale bars= 10 m. (D) Quantification of the experiment in (C) showing the number of BRCA1 (left panel) and RAD51 (right panel), respectively, nucleolar caps (n= 100 cells). Graphs represent a single replicate (experiment was performed in triplicate), and bars represent mean.

**Figure 5–figure supplement 1**

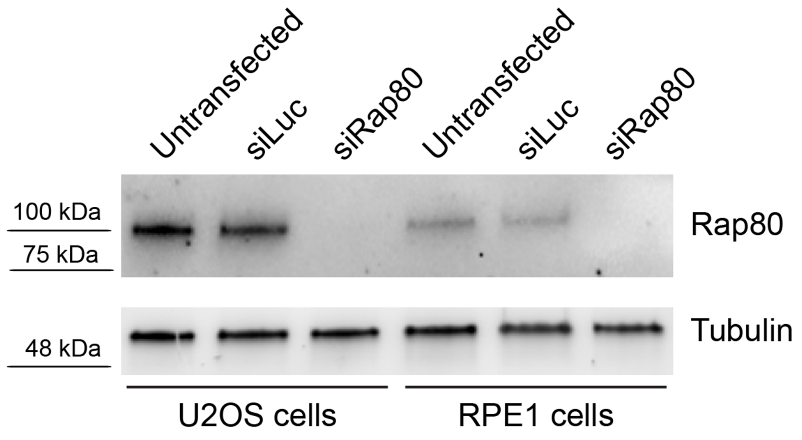


Western blotting of extracts from U2OS and RPE1 cells, either untransfected or transfected with siLuc and siRAP80, respectively.

### Figure 6–figure supplement 1

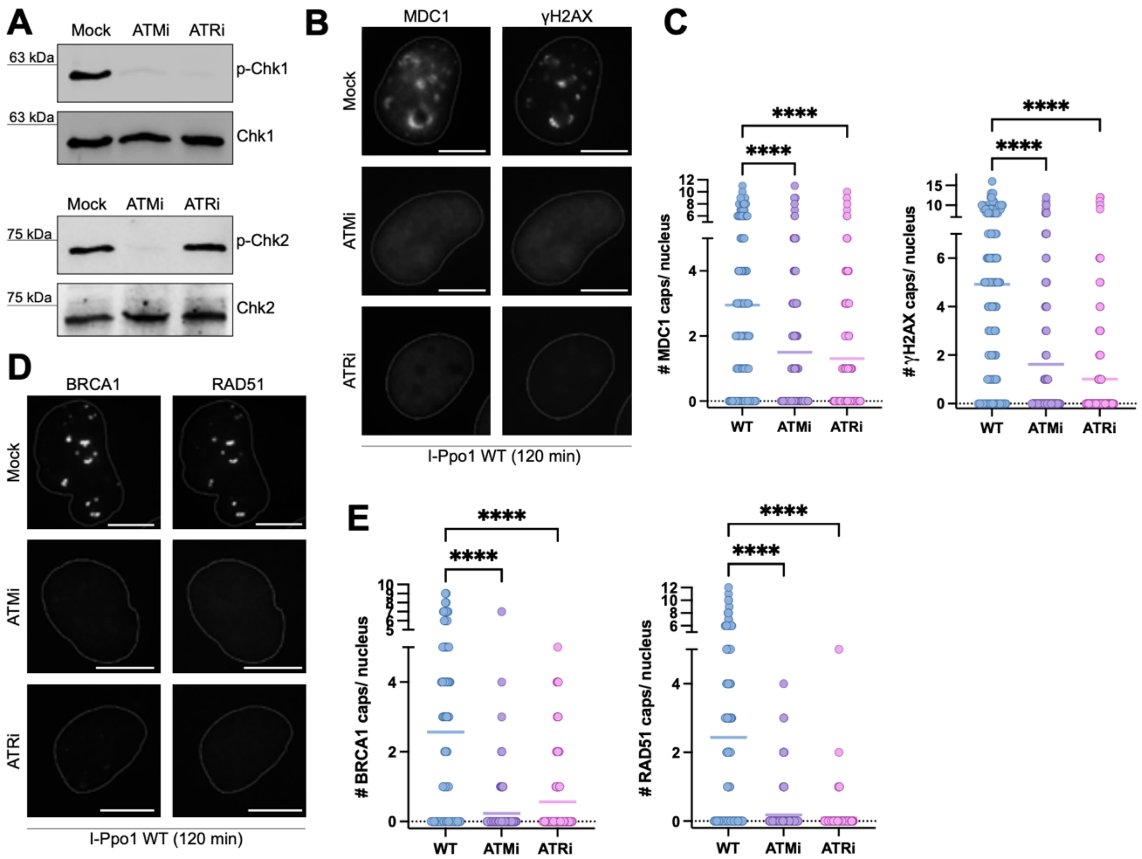


(A) Western blotting of extracts from U2OS cells treated with or without ATM and ATR inhibitors and transfected with I-Ppo1 WT for 2h. (B) Localization of MDC1 and γH2AX after 2h of I-Ppo1 WT transfection in U2OS cells treated with or without ATM or ATR inhibitors. (C) Quantification of the experiment in (B) showing the number of MDC1 (left panel) or γH2AX (right panel) nucleolar caps per nucleus (n = 100 cells). Graphs represent a single replicate (experiment was performed in duplicates) and bars represent mean. (D) Localization of BRCA1 and RAD51 after 2h of I-Ppo1 WT transfection in U2OS cells treated with or without ATM or ATR inhibitors. All scale bars= 10 $\mu$m. (E) Quantification of the experiment in (D) showing the number of BRCA1 (left panel) or RAD51 (right panel) nucleolar caps per nucleus (n= 100 cells). Graphs represent a single replicate (experiment was performed in duplicates) and bars represent mean.

### Figure 7–figure supplement 1

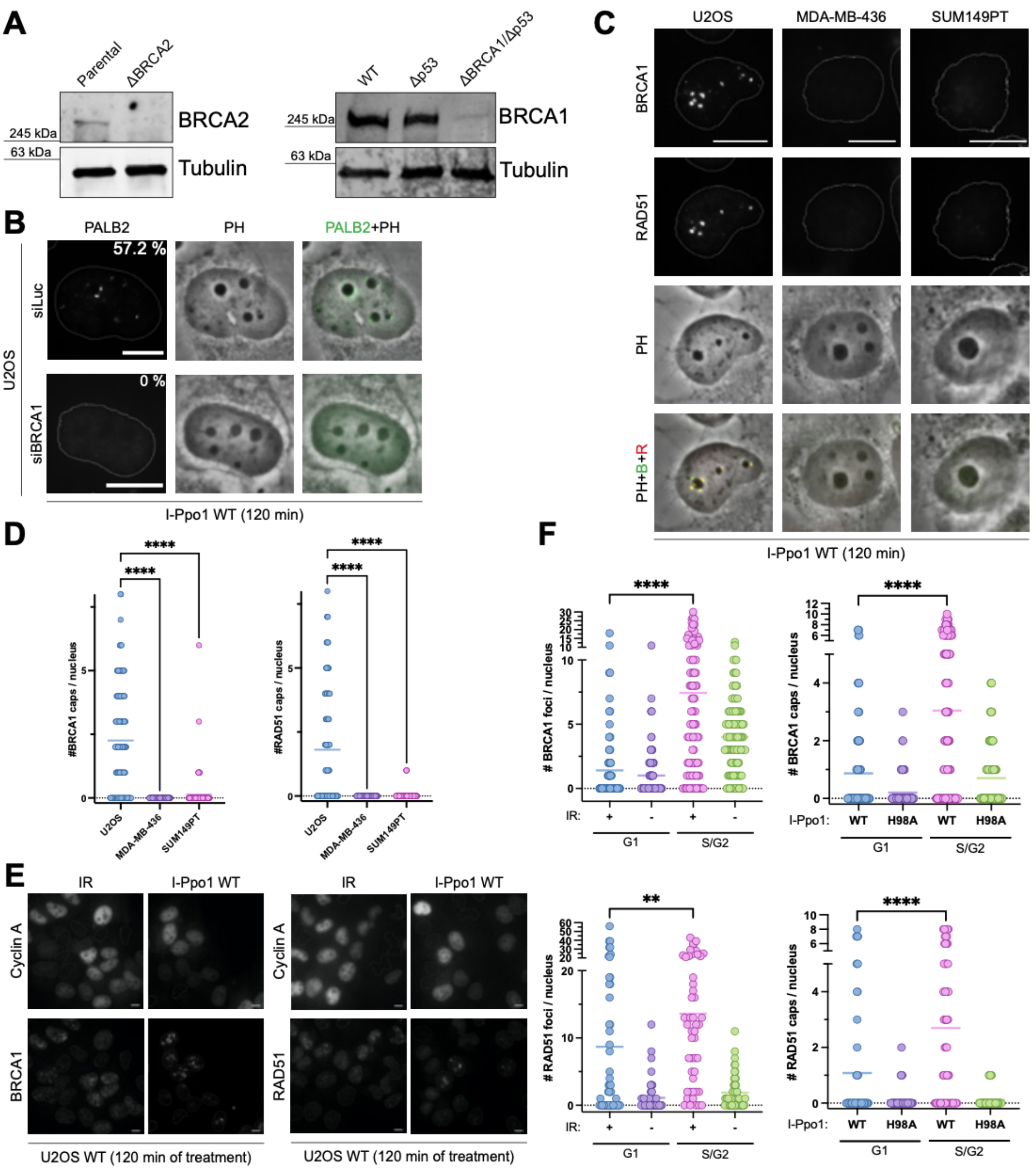

(A) Western blotting of extracts from parental and ΔBRCA2 DLD1 cells and parental and ΔBRCA1 ΔTP53 RPE1 cells, respectively. (B) Localization of PALB2 after 2h of I-Ppo1 WT transfection in U2OS cells treated with siLuc or siBRCA1. Percentage shows the number of PALB2 positive cells in the population (n= 110 cells). All scale bars= 10 $\mu$m. (C) Localisation of BRCA1 and RAD51 in U2OS WT, MDA-MB-436, and SUM149PT cells after 2h of I-Ppo1 WT transfection. All scale bars= 10 $\mu$m. (D) Quantification of the experiment in (C) showing the number of BRCA1 (left panel) or RAD51 (right panel) caps per nucleus (n= 100 cells). Graphs represent a single replicate (experiment was performed in duplicate) and bars represent mean. (E) Localisation of BRCA1 and RAD51 after irradiation or I-Ppo1 WT transfection in U2OS cells. (F) Quantification of the experiment (E) showing the number of BRCA1 foci (first panel) or the number of BRCA1 nucleolar caps (second panel) per nucleus (n= 100), and the number of RAD51 foci (third panel) or the number of RAD51 nucleolar caps (fourth panel) per nucleus (n= 100) in U2OS cells treated with irradiation or I-Ppo1 WT, respectively. Graphs represent a single replicate (experiment was performed in duplicate), and bars represent mean.
